# A centrin–Sfi1 myoneme fishnet powers ultrafast calcium-triggered contraction in the giant ciliate *Spirostomum ambiguum*

**DOI:** 10.1101/2024.11.07.622534

**Authors:** Joseph Lannan, Carlos Floyd, L. X. Xu, Peter M. Thompson, Connie Yan, Wallace F. Marshall, Surirayanarayanan Vaikuntanathan, Aaron R. Dinner, Jerry E. Honts, Saad Bhamla, Mary Williard Elting

**Affiliations:** Physics, North Carolina State University; Chemistry, University of Chicago; Chemical and Biomolecular Engineering, Georgia Institute of Technology; Molecular and Structural Biochemistry, North Carolina State University; Molecular Education, Technology, and Research Innovation Center, North Carolina State University; Biochemistry and Biophysics, University of California San Francisco; Biology, Drake University; BioFrontiers Institute and Department of Chemical and Biological Engineering, CU Boulder, Colorado, USA; Cluster for Quantitative and Computational Developmental Biology, North Carolina State University

## Abstract

*Spirostomum* is a giant unicellular ciliate that contracts to a quarter of its body length in less than five milliseconds, achieving an order of magnitude higher fractional shortening rate than actomyosin-based systems. This ultrafast contraction is powered by myonemes, calcium-activated protein networks at the cortex whose biochemical mechanism remain unclear. We quantify changes in cortical microtubules, membrane ruffles, and the fishnet-like myoneme mesh during contraction, and develop multiscale models that connect local myoneme shortening to whole-cell shape change. Centrin and an Sfi1 homolog co-localize with the myoneme by immunofluorescence and localize to the myoneme by immunogold electron microscopy. Coarse-grained mesh simulations reproduce the measured deformations and show that fishnet geometry, together with volume conservation, leads to uniform contraction. Finally, we reconstitute a *Spirostomum* centrin–Sfi1 repeat complex *in vitro* and measure calcium-dependent compaction and self-association, supporting a molecular basis for myoneme contractility. Together, these results support a multiscale model in which calcium-responsive centrin– Sfi1 structures are the central contractile element in *Spirostomum* and suggest design principles for fast, calcium-triggered, chemomechanical contractile networks that operate without actomyosin or ATP.

**SIGNIFICANCE STATEMENT:** Many cells change shape using actomyosin, but some protists contract using calcium-activated protein networks called myonemes. We combine quantitative imaging, electron microscopy, multiscale modeling, and *in vitro* reconstitution to link molecular-scale mechanisms to the millisecond shortening of the giant ciliate *Spirostomum*. Centrin and an Sfi1 homolog co-localize in a fishnet-like cortical mesh, and simulations show that this geometry can reproduce the observed whole-cell shape change under volume conservation. Purified centrin–Sfi1 complexes undergo calcium-dependent compaction and self-association, supporting a protein-scale switch that can drive myoneme contraction. These results connect calcium signaling to whole-cell mechanics and suggest principles for designing fast, ATP-independent bioinspired actuators and synthetic cellular machinery.

## INTRODUCTION

The giant unicellular ciliate *Spirostomum* has the ability to rapidly contract from a length of ~1 mm to ~300 *µ*m in as little as 5 ms [1], a speed upwards of ~100 body lengths per second. In contrast, individual muscle fibers are similar in size and can shorten by a similar fraction [2], but their typical maximum velocity is approximately 10-fold slower [3]. Although related ciliates, such as *Stentor*, also exhibit large scale contractions, likely with biochemically similar machinery, *Spirostomum* is notable for its speed, especially relative to its size [4– Additionally, *Spirostomum* is able to quickly repeat this motion, resetting within a few seconds, unlike other fast but one-shot biological firing mechanisms such as nematocysts in jellyfish tentacles [7]. Interestingly, this contraction appears to be powered by a molecular mechanism distinct from conventional cytoskeletal filaments such as actin and microtubules and is triggered by calcium ions, without direct association with ATP or GTP [8, 9]. Thus, *Spirostomum* offers a unique system for testing the physical limits of biological components to generate power, from the molecular to the organismal level. Ultimately, understanding this machinery may yield applications in the generation of mechanically robust and re-configurable cytoskeletal structures.

The unconventional cytoskeletal structures that drive the contraction of *Spirostomum* and related ciliates are called myonemes. Myonemes are long fibrous networks in the cortex that contract in response to calcium [10–12] and are thought to be responsible for cell shortening [13, 14]. While they are far from fully characterized, previous observations of myonemes by electron microscopy showed fibrous bundles that change appearance and increase in density under contraction [15–17].

Although actomyosin powers many biological contractions, it has not been observed in the myoneme. Instead, myonemes are rich in centrin and large homologs of its binding partner Sfi1 [5, 18, 19]. The presence of centrin and Sfi1 in myonemes is as surprising as the absence of actomyosin, since homologous proteins in other systems have very different known functions. In *S. cerevisiae*, centrin/Sfi1 filaments form a largely linear structure that may act as a molecular ruler in constructing the *S. cerevisiae* spindle pole body [20, 21] There is precedent for calcium-triggered contraction of centrin-based structures in algal striated flagellar roots, where centrin was first discovered [22]; but despite speculation based on this earlier evidence [23], *S. cerevisiae* centrin/Sfi1 filaments show no evidence of kinking, bending, or sliding, even in the presence of millimolar calcium ion concentrations [20]. Thus, the molecular mechanism by which ciliate centrin and Sfi1 couple calcium influx to force generation remains unclear.

Alongside myonemes, microtubules have been identified as a key component of the *Spirostomum* cortical cytoskeleton, although their potential role in contraction and/or re-elongation is not yet clear. Cortical microtubules decorate the surface of *Spirostomum* and form interconnected bundles along rows of basal bodies [24]. Previous models have proposed that a physical link between myonemes and microtubules could allow them to transmit forces without slipping past each other [16, 25], or that the antagonistic action of myonemes and microtubules might support repeatable contraction-elongation cycles [13, 14]. Yet the interactions between microtubules and myonemes and their mechanical contributions to changes in organismal shape remain largely unresolved.

A challenge in achieving a complete understanding of *Spirostomum* contraction is the range of length, time, and force scales that must be described. We have recently measured the contraction dynamics and used them to model in one dimension how the onset of contraction propagates along the entire organism [26]. However, a complete model of *Spirostomum* contraction would describe not only the dynamics of triggering but also how molecular events generate and direct force to induce appropriate changes in three-dimensional shape. Here, we build toward such a model with an approach that spans scales, from the molecular to the organismal. We quantitatively characterize the rearrangements within the myoneme and of the organism as a whole by light and electron microscopy. These data, in turn, inform computational models of contraction on both a coarse-grained and molecular scale. Additionally, we used *in vitro* reconstitution of recombinant proteins to directly demonstrate the contractile ability of *Spirostomum* centrin/Sfi1. These results led us to propose a multi-scale mechanism explaining centrin/Sfi1 force generation.

## RESULTS

### Immunofluorescence microscopy reveals organismal-scale changes in *Spirostomum* cortical structure under contraction

The structural changes that accompany contraction can provide information on force generation and the subsequent storage and dissipation of mechanical energy. By altering fixation conditions [27, 28] (see Materials and Methods), we can preserve cells in elongated (length = 884 ± 111 µm, N=7) or contracted (length = 306 ± 32 µm, N=12) states. This three-fold average decrease in length is accompanied by an increase in diameter from 89 ± 10 µm (N=7) to 117 ± 11 µm (N=12). Using immunofluorescence microscopy, we visualize the accompanying changes in microtubules, membrane, and centrin (Fig. 1A). Each of these structures exhibits significant but distinct rearrangements under contraction.

**FIG. 1:**
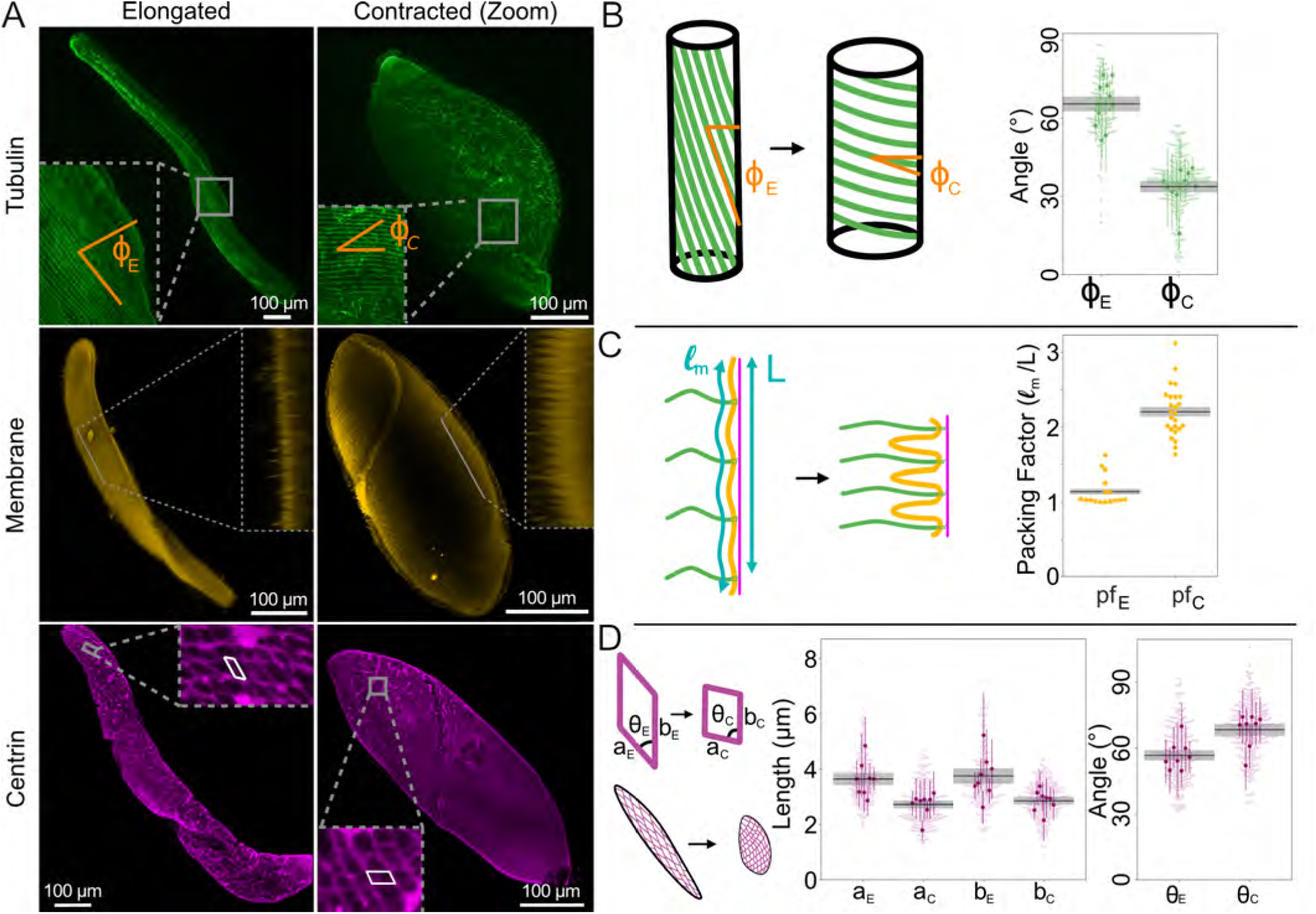
Changes in *Spirostomum* cortex in elongated (subscript E) vs. contracted (subscript C) cells quantified by immunofluorescence. **A** Representative example maximum intensity projections of confocal fluorescence microscopy z-stacks of *Spirostomum* stained via TAP952 (anti-tubulin, green, top), Cellmask Orange (membrane, yellow, middle), and 20H5 (anti-centrin, magenta, bottom). Membrane insets show single slices projected in the perpendicular plane. For space, elongated and contracted cells are shown at different scales. **B**–**D** Quantification of image features, as shown in cartoons. Black lines and gray bars show mean and standard error of the mean (SEM) over N=number of cells in each measurement. Larger and darker markers in **B** and **D** show means from individual cells with standard deviation (as colored lines) and lighter, smaller markers show swarm plots of all individual measurements, measurements in **C** are per individual cell. **B** Microtubule pitch angle, measured by approximating the centerline of the organism and taking the complementary angle between the microtubule and the center line ϕ, N = 10 cells for each condition. **C** Packing factor due to membrane buckling, L is the end-to-end distance and *l*_*m*_ is the length along the membrane, N=10 cells for each condition, measured from orthogonal re-projection along a straight section of the organism. **D** Length and angle of myoneme mesh segments, N=8 cells for each condition, 30 segments (a and b each) measured evenly split between each side and the center of organism.

Cortical microtubules form long helical structures that change pitch under contraction (Fig. 1B). Quantification shows that the mean pitch angle (ϕ) when elongated is 64°± 3°, and when contracted, it is 34°± 2° (SEM reported on N = 10 cells, p = 1 · 10^−7^). However, due to the simultaneous change in diameter of the organism, these bundles undergo very little change in radius of curvature despite their change in pitch (Fig. S1). In fact, we estimate that, unlike the contribution of microtubule bending to neck elongation recently reported in *Lacrymaria olor* [29], microtubule bending is unlikely to make a significant contribution either in opposing contraction or powering elongation in *Spirostomum* (see Supplementary Discussion 1 and Fig. S1).

Next, we visualize the membrane organization of elongated and contracted cells using CellMask Orange plasma membrane stain (Fig. 1A). By collecting z-stacks, we observe that the membrane buckles and forms ridges under contraction (Fig. 1C). We quantify these ridges by visualizing them in cross-section (Fig. 1A, inset) and calculating their packing factor (*pf*), defined as the length of the contoured surface divided by the end-to-end distance. We measured *pf* = 1.14 ± 0.03 in elongated *Spirostomum* and 2.20 ± 0.07 in contracted *Spirostomum* (SEM reported on N = 10 cells, p = 3 · 10^−15^), indicating that there is twice as much membrane per length in contracted cells compared to elongated cells. This measured packing factor is consistent with maintaining the total membrane surface area under contraction, since ridges allow the effective surface area of the organism to decrease by effectively storing membrane as the organism contracts. We estimate the energy stored in membrane bending during contraction (see Supplementary Discussion 1) and find that it is ~ 100 fJ, so small as to contribute negligibly to the mechanics of the system as a whole.

Finally, we visualize the myoneme architecture in both contracted and elongated cells by staining with the centrin antibody 20H5 (Fig. 1A, S2). In qualitative agreement with previous observations [5, 30], we observe a mesh of packed, parallelogram-shaped bundles that tile the entire surface in both elongated and contracted cells (Fig. 1A, D). Quantification of their dimensions (Fig. 1D) reveals a decrease between elongated and contracted cells in the lateral direction (*a*) of 30%, from 3.6±0.2 µm to 2.8 ± 0.1 µm (*p* = 0.007), and in their longitudinal direction (*b*) of 24%, from 3.7 ± 0.3 µm to 2.8 ± 0.1 µm (*p* = 0.01) (SEM reported over N=8 cells). Accompanying the myoneme shortening, we observe shearing of each individual parallelogram, with an increase in the angle of the vertex θ from 58.8° ± 2.9° to 68.4° ± 2.8°(*p* = 0.03) (SEM reported on N = 8 cells). As previously demonstrated [5], we confirm that centrin and Sfi1 colocalize in the myoneme (Fig. S3). Using the geometry we measure here and scaling previous measurements of the total contractile force at the organismal-scale [11], we estimate that each unit of myoneme (i.e., each bundle that comprises the side of a parallelogram) must generate ~ 1000 pN of force during contraction (Supplementary Discussion 1).

### Coarse-grained mesh models demonstrate how myoneme bundle organization supports macro-scale contraction of the organism in 3D

To construct a mechanical description of *Spirostomum* contraction at the organismal scale, we developed a coarse-grained mesh model of the cortical cytoskeleton (Fig. 2). This model helps us understand the mechanical contributions of the cytoskeletal elements observed by immunofluorescence (Fig. 1). We modeled the myoneme system as a quadrilateral mesh that encloses a rounded cylinder (Fig. 2A). This allows us to capture the contributions of volume conservation and the overall shape change of the organism. We treat the edges of the quadrilateral unit cell as harmonic springs whose rest lengths shrink by a factor (*γ*) during contraction. To simulate contraction, we numerically minimize the total energy objective function with respect to the configuration of the mesh under the new set of rest lengths. Since previous measurements of surrounding flows show no evidence of significant fluid emission during contraction [1], we assume the conservation of total volume. To enforce this constraint, we triangulate the surface [31] by dividing each quadrangular unit cell of the mesh into two and augmenting the energetic objective function to include a stiff quadratic penalty against deviating from the original volume. To represent the possible mechanical contribution of microtubules, we also include an optional torsional spring term that penalizes the untwisting of adjacent layers; we study whether this term is necessary to obtain the contracted shapes observed *in vivo*. The total energy function thus represents the balance of local elastic penalties (on the myoneme strands and microtubules) with the strict global constraint of zero volume change. See Materials and Methods for additional details and Supplementary Table S2 for model parameters.

**FIG. 2:**
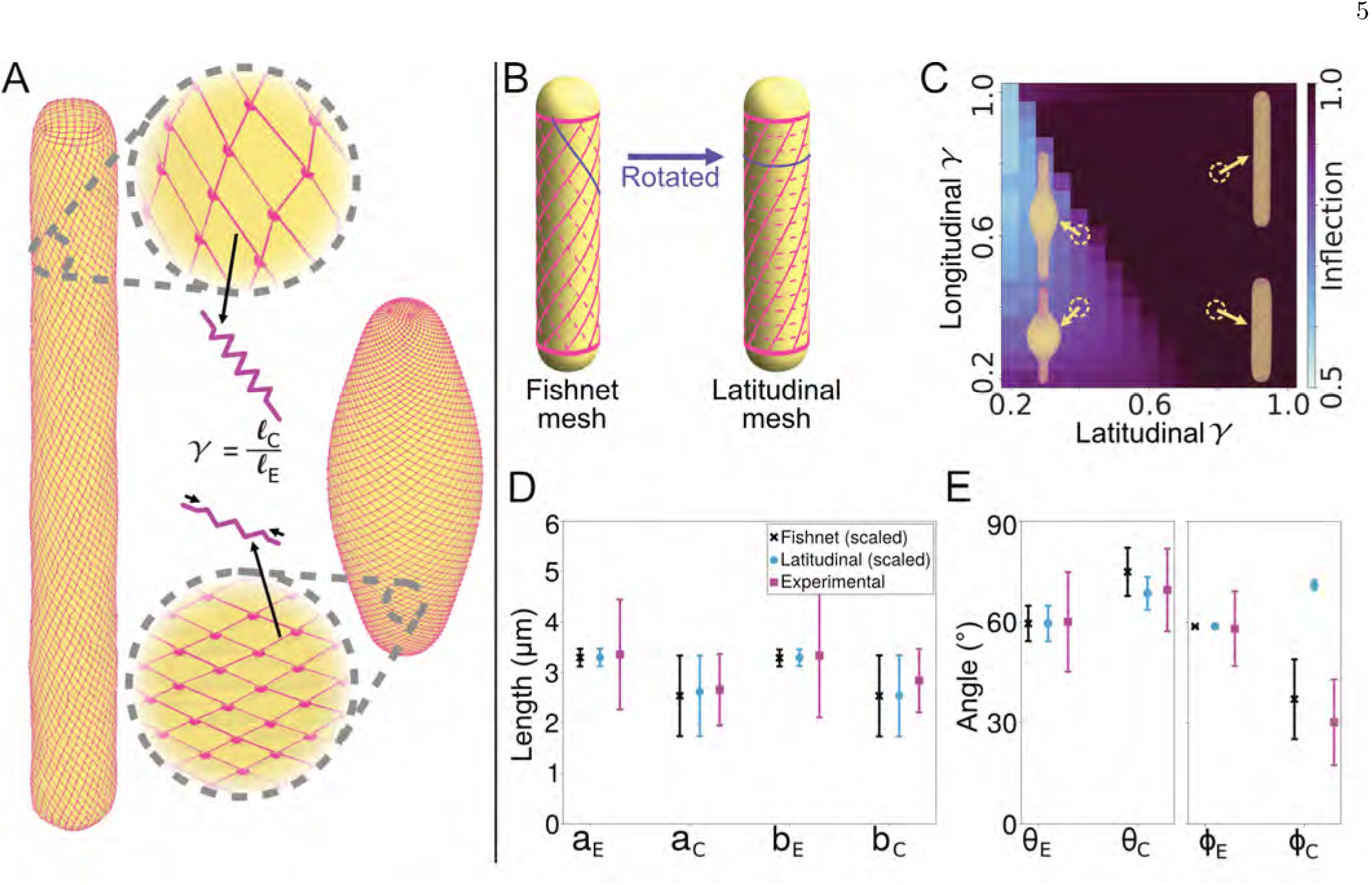
Coarse-grained fishnet mesh model of the *Spirostomum* cortex recapitulates shape changes observed *in vivo*. **A** Cartoon of model, which represents myonemes as a quadrilateral mesh of springs (magenta) that surrounds a rounded cylinder (yellow). We simulate contraction by shrinking the rest lengths of the springs by a fraction *γ* and minimizing the total energy. **B** Schematic illustration of the structure of the fishnet (left) and latitudinal (right) mesh. The latitudinal mesh structure can be obtained from the fishnet mesh by rotating the dashed lines to connect opposing vertices. **C** Heatmap of the inflection metric (see Supplementary Discussion 2 for definition) for the latitudinal mesh with varying shrinking factor *γ* for both edges of the mesh. Example structures are displayed near their corresponding locations on the heatmap. **D** and **E** Comparison of simulated and experimental measurements before and after contraction. Error bars represent standard deviation between individual filaments across the model and all samples from 1D. **D** Edge lengths of the unit cells, a and b are the edges of the parallelograms that make up the mesh, subscript E: elongated, subscript C: contracted (cf. Fig. 1**D**). The coarse-grained simulation unit cells are scaled by a constant factor to allow comparison. **E** Angle of the unit cell parallelograms θ (cf. Fig. 1**D**) and helix twisting angle ϕ (cf. Fig. 1**B**).

To test the importance of the observed myoneme geometry for effective contraction, we consider two types of mesh structures: “fishnet” and “latitudinal.” In the fishnet mesh, which corresponds to the geometry found *in vivo*, sets of intersecting myoneme strands run along the length of the cylinder as helices with opposite handedness to each other (Fig. 2B). The initial helix angles of these strands are set to coincide with measurements of the elongated microtubule helix angles (Fig. 1B) and the angles of the myoneme unit cells (Fig. 1D). In the latitudinal mesh, which we consider as a hypothetical alternative mesh geometry, we rotate one set of strands to make it perpendicular to the long axis (dashed lines in Fig. 2B), thereby forming closed loops around the cylinder’s “lines of latitude.”

We find that the latitudinal mesh requires fine-tuning and, even with a best-fit set of parameters that includes torsional resistance, does not satisfactorily reproduce the experimentally measured changes in the mesh structure (Fig. 2C-E and Fig. S4). By contrast, we find that the contraction of the fishnet mesh structure quantitatively reproduces the experimental measurements of the changes in the side lengths of the unit cells (Fig. 1D and 2D), the angles of the unit cells (Fig. 1D and 2E), and the angle of the helix (Fig. 1B and 2E). Interestingly, we find that this agreement occurs without any torsional resistance of the microtubules. We observe that agreement occurs fairly robustly over a range of shrinking factors for the two sets of myoneme strands. Overall, the model suggests that the fishnet mesh structure is a key determinant of the experimentally observed contraction, which can explain the changes in the unit cell structure and helix angle.

### TEM shows mesoscale changes in myoneme bundles from a loose network to a dense contracted structure

To examine the underlying structural changes in the myoneme that facilitate contraction, we performed transmission electron microscopy (TEM) on *Spirostomum* treated with and without EGTA, which preserves it in an elongated or contracted state, respectively (Fig. 3). Comparisons between these two states provide insight into the molecular and structural bases of contraction across length scales. We also performed immuno-gold labeling to verify the identity of myonemes in the TEM and to determine the localization of centrin and Sfi1 in the myoneme at this scale.

**FIG. 3:**
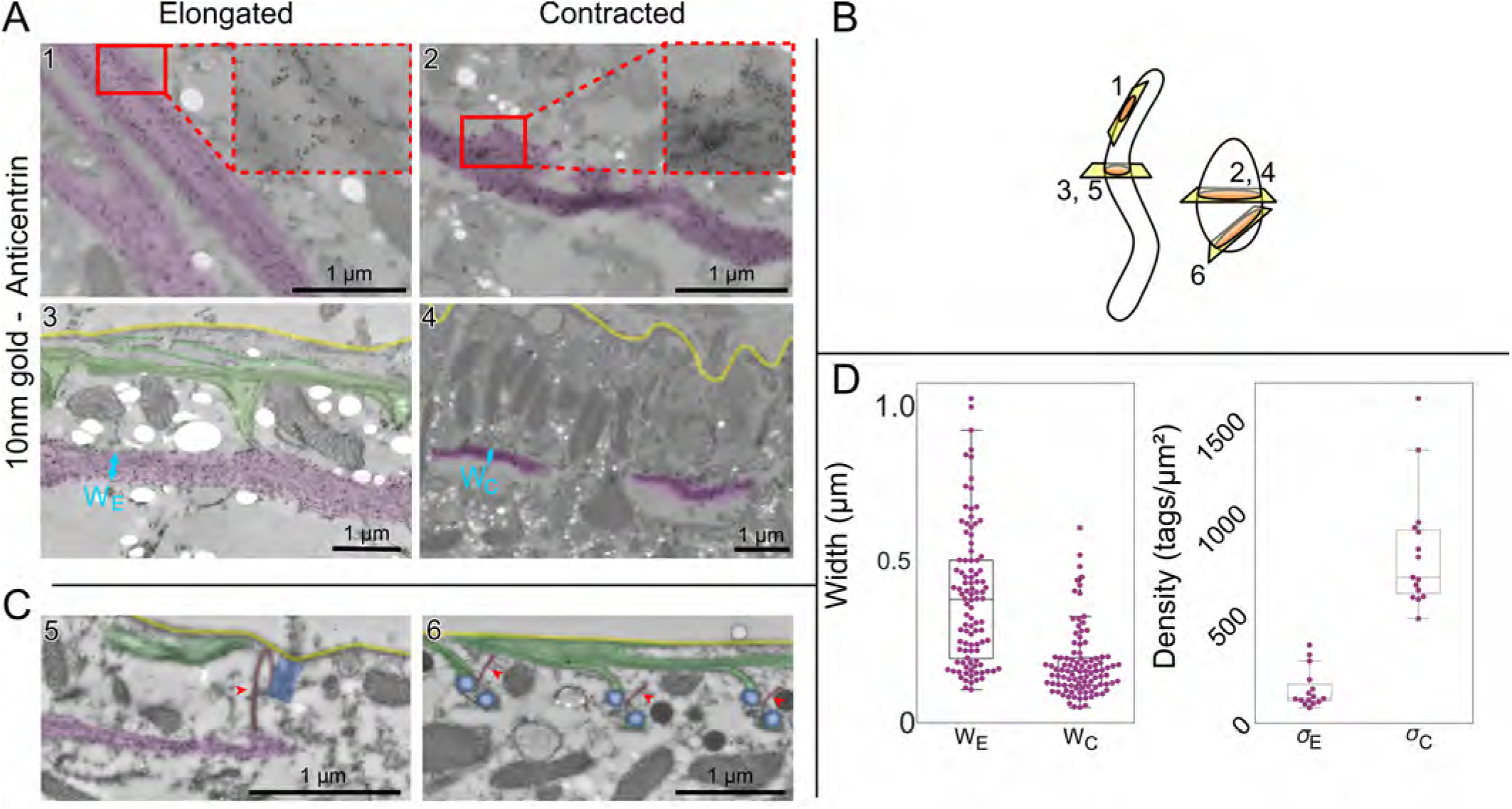
Immunogold TEM reveals changes in centrin-dense myoneme filaments. Images highlighted for myoneme bundles (magenta), plasma membrane (yellow) and microtubules (green). Highlight colors are hand annotated. **A** Representative example images of immunogold labeled with 20H5 anti-centrin. In elongated cells (left, images 1 and 3), myoneme bundles appear as a loose meshwork with some long connected filaments, while in contracted cells (right, images 2 and 4), bundles appear as dense structures with no discernible fine details. There are holes present in some samples (bright white ovals) due to the fragility of the uncoated grids used to increase labeling efficiency. **B** Cartoon showing approximate direction of cuts. **C** Myoneme and microtubule cytoskeletons connect near basal bodies (highlighted in blue). Left (elongated cell), example image showing their fiber connection (red arrows and red highlights). Right (contracted cell), A perspective perpendicular to the axis through the center of the basal bodies and perpendicular to 5. This view shows a similar fiber (red arrowhead) that likely connects to the myoneme in a different plane. Note that image 5 had reduced labeling efficiency as it was performed on a backed grid. It was included due to the rarity of obtaining a section that depicts the full fiber. **D** Quantification of myoneme bundle width (left) measured perpendicular to the cell membrane at regular increments across each individual filament to show variation in a single filament. Images used were perpendicular to the membrane e.g. 3 and 4 where branching was not observed (elongated: N=4 images with 3 preparation repeats, contracted N=5 images with 3 preparations) and tag density (right; 15 measurements shown, 3 repetitions of the preparation, measurement is average over a bundle). Box plot center line, median; box limits, upper and lower quartiles; whiskers, 1.5x interquartile range; points, outliers. Each preparation contained multiple cells; it is unknown whether the images in a single preparation were of different cells.

In both elongated and contracted cells, the centrin antibody (20H5) densely and specifically labels the myoneme, as well as the centrin located in the basal bodies and some additional cortical structures (Fig. 3A, B). In elongated cells, we observe a loose network with filaments that run through the myoneme (Fig. 3A). It is unclear whether some of the additional cortical staining represents other centrin-containing structures or non-specific binding of the 20H5 antibody. In contracted cells, the bundles are highly dense, and we cannot resolve their substructure in the TEM images (Fig. 3A), consistent with previous observations [17]. In both elongated and contracted cells, the centrin label is roughly uniform throughout the entire myoneme, verifying it as a core myoneme component (Fig. 3 A). To label Sfi1, we used peptide antibodies raised against sequences from the related ciliate *Stentor* (see Materials and Methods and [19]). Perhaps unsurprisingly, these Sfi1 antibodies had a lower affinity in *Spirostomum* but still sparsely labeled myonemes, as expected from our own observations (Fig. S3) and previous immunofluorescence studies [5].

In both elongated and contracted cells, we observe associations between microtubules and myonemes at basal bodies through slices that happen to intersect with the fiber (Fig. 3C). This direct interaction between myonemes and basal bodies has been speculated [5] and observed in a previous report [17]. Here, we find that cortical microtubule bundles are also involved and that these interacting microtubules often show immunogold labeling for centrin, suggesting that centrin may play a role not only in filament structure but also in anchoring myonemes to other cortical structures to distribute force and maintain organization under contraction (Fig. 3A, panel 3).

We quantified the width of the bundles in elongated and contracted states, finding a decrease by a factor of 3 (Fig. 3D), from a width of 0.61 ± 0.06 µm in elongated bundles to 0.20 ± 0.03 µm in contracted bundles (*P* = .004, SEM calculated with N=3 replicates each). Furthermore, we quantified the density of immunogold tags, observing an increase in tag density by a factor of 8 (Fig. 3D), from 116 ± 23 tags/µm^2^ in elongated cells to 913 ±154 tags/µm^2^ in contracted cells (*P* = .04, SEM calculated with N=3 replicates each). Thus, the structural changes we observe at the nanoscale are consistent with the interpretation that myoneme rearrangements drive the contraction of the organism as a whole.

### Skeletonization analysis of TEM reveals the micro-scale organization of the elongated myoneme

The mesh-like structures that we observe within the myoneme bundles of EGTA-treated elongated *Spiros-tomum* show some regularity, which we characterize through semi-automated skeletonization analysis (Fig. 4A). We then process the skeletonized images to identify the intersections and branches (Fig. 4B). While significant variations, likely due to filaments coming in and out of the section, limit the precision of the skeletonization, we quantify that the primary angle of the junctions is ≈ 100 − 135° and most junctions include 3 connections, consistent with an approximately hexagonal mesh (Fig. 4C). Among this branching structure, we qualitatively observe many consistent, long strands traversing the long axis of the myoneme bundle. Although this structure is somewhat different from that previously described by Ishida et al. [16], our preparation involved a higher concentration of EGTA for a shorter time, which may have caused the myoneme to access a more relaxed state. These long strands may be individual *Spirostomum* Sfi1 (GSBP1/2) filaments, which are predicted to be up to microns long [5].

**FIG. 4:**
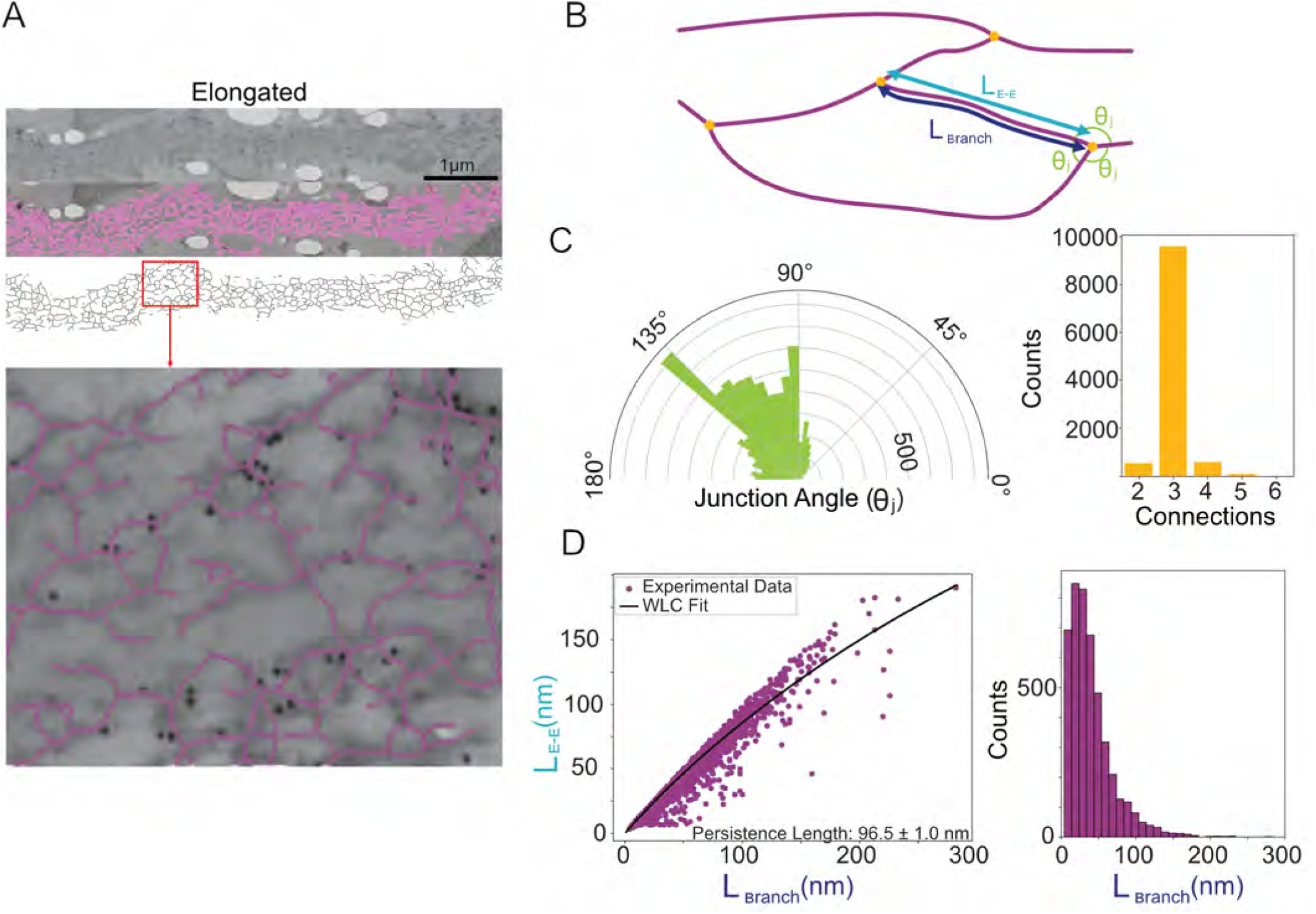
Skeletonization analysis reveals structural features of elongated myonemes. **A** Selected image showing segmentation analysis (magenta) and skeletonization. Inset shows overlay of original image with skeletonized data. **B** Cartoon showing features quantified from the skeletonization. Branches (magenta) of segment lengths *L*_*branch*_ and end-to-end distances *L*_*E*_−_*E*_ meet at intersections (yellow dots) with junction angles θ_*j*_. **C** Polar histogram of junction angles (left) and number of branches at each intersection (right) (N=5 images, 10,816 total measurements). **D** Left, plot of branch length versus end-to-end length, fit to a worm-like chain (WLC) model (see Methods) to estimate the persistence length. Right, histogram of branch lengths (excluding dead end branches), demonstrating notable variation.

By comparing the distribution of lengths of the skeletonized filaments in each branch with their end-to-end lengths, we estimate their persistence length (Fig. 4D). Assuming that there is little external force on the elongated myoneme, we apply a worm-like chain model (WLC), where the persistence length *P* is defined as the characteristic length over which the direction of the filament becomes uncorrelated with itself, i.e., 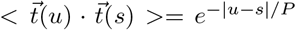, where t is the tangent vector along a polymer, u and s are positions along the polymer [32]. This model produces a fitted persistence length of 96 ±1 nm (SEM calculated from the covariant matrix reported by the scipy python package [33]). Consistent with the hypothesis that these filaments may represent individual centrin-bound Sfi1 helices, this estimate is similar to measurements of the calmodulin-stabilized lever arm of myosin V, which has a persistence length of ≈ 150 nm [34] and shares a gross structural similarity with centrin-bound Sfi1.

### Molecular structural predictions suggest potential nano-scale mechanisms of myoneme contraction

Although the complete biochemical composition of the myoneme is not fully established, the two known components in both *Spirostomum* and related systems are centrin and homologs of Sfi1 [5, 35, 36]. In *Saccharomyces cerevisiae*, the long alpha-helical protein Sfi1 (molecular weight 110 kDa) forms a scaffold that includes up to 24 centrin binding sites, each approximately 23-36 amino acids long (Supplementary Table S4), creating filamentous structures mediated by centrin-centrin interactions [20, 37]. To gain insight into how homologous components might have been adapted to provide a molecular basis for contraction-generating myoneme filaments in *Spirostomum ambiguum*, we first analyzed their sequences and modeled their structures using AlphaFold (Fig. 5A and Supplementary Fig. S5) [38–40]).

**FIG. 5:**
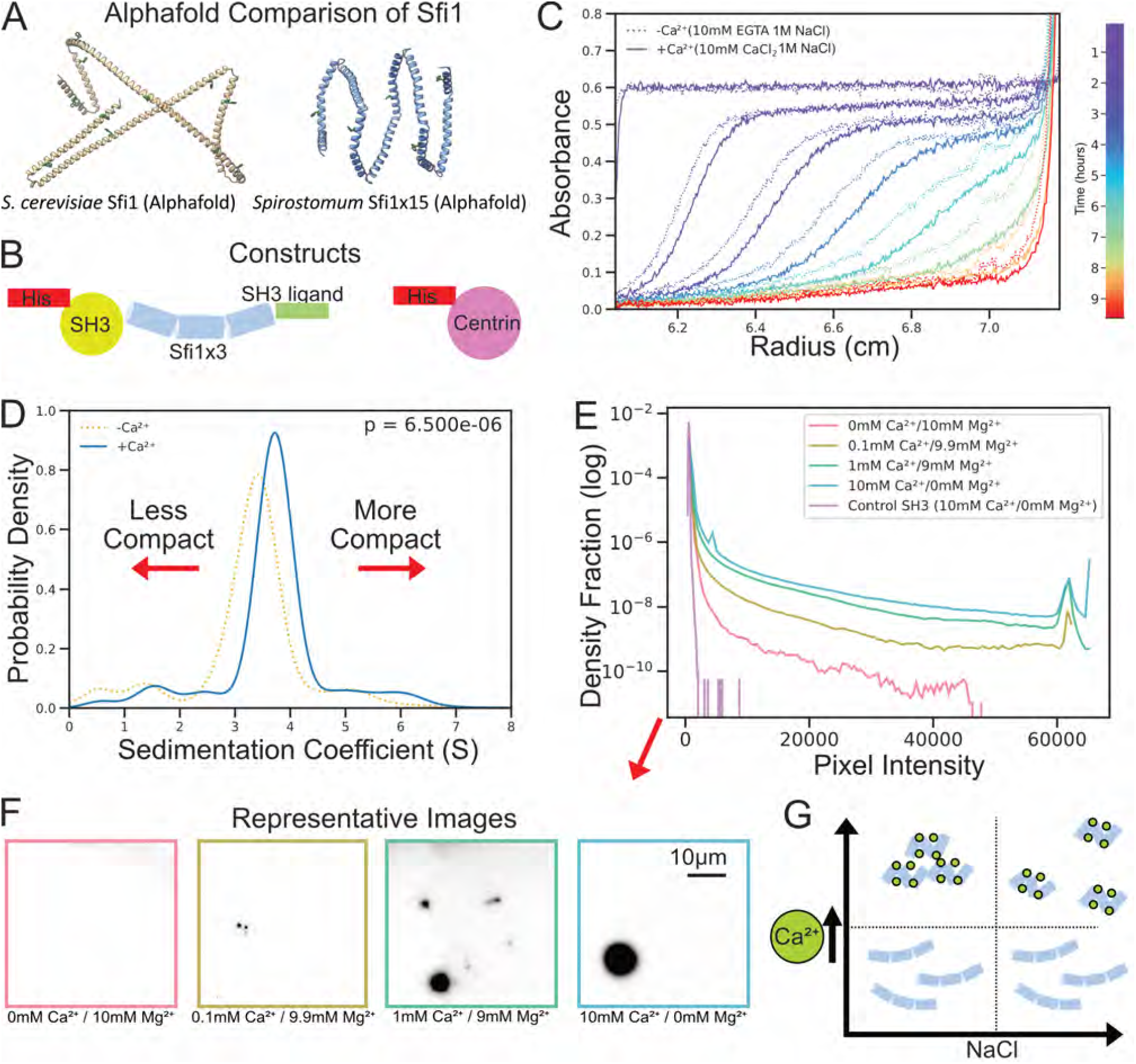
Centrin Sfi1 constructs contract and aggregate with calcium. **A** Comparison of *S. ceversiae* and *Spirostomum* Sfi1 AlphaFold models showing the helix breaking residues in *Spirostomum* Sfi1 repeats cause increased kinking and compaction. **B** Diagrams of the protein constructs used in subsequent experiments to test the theory that the helix breaking residues allow for bundle compaction with the addition of centrin. **C** Example of raw analytical ultracentrifuge traces, with multiple time points over a run overlaid in different colors. Scans were acquired sequentially; to avoid artifactual offsets from scan timing, the –Ca^2+^ condition was selected after the +Ca^2+^. The construct used was a 3x 69 amino acid repeat of Sfi1 co-purified with Centrin. These samples were run in high salinity which prevented larger scale aggregation of the construct with calcium. **D** Fit for sedimentation coefficient, averaged over 6 runs for the AUC data shown in **C**. T-test shown is for the mean of the 6 runs. **E** Histogram of epi-fluorescence images of Sfi1×3 + centrin constructs, with increasing calcium conditions showing increased aggregation with the addition of calcium. Magnesium was used as a divalent cation control showing specificity for calcium. Two independent purification; two samples per purification; 100 fields per sample; pooled analysis. **F** Selected images from **E** showing large scale aggregation of the construct with the addition of calcium. With constant concentration of total divalent cations (Ca^2+^ and Mg^2+^), an increase in calcium tends towards larger scale aggregation as shown by the larger and darker spots in the selected images. **G** Proposed phase diagram of Sfi1/centrin response to calcium and NaCl conditions. Increases in calcium cause individual centrin/Sfi1 complexes to compact and aggregate. Aggregation tends to be inhibited by an increase in NaCl concentration.

*Spirostomum ambiguum* cells were processed to isolate cortical cytoskeletons and enrich myonemal proteins. The proteins in these preparations were separated by gel electrophoresis, and protein bands were excised for mass spectrometric analysis. We identified putative centrin and Sfi1 protein sequences derived from *Spirostomum ambiguum* transcriptomics data deposited in NCBI (see Materials and Methods, Supplementary Tables S3, and Supplementary Data). We found several proteins that contain multiple tandem repeats of a 69 amino acid sequence similar to Sfi1-like centrin-binding proteins in *Paramecium tetraurelia*, as well as 2 centrins that are also consistent with the mass spectrometry of the isolated myoneme. Although the reads we identified from the deposited transcriptomics are relatively short, the 69 amino acid repeats still show close homology to the Giant Spasmoneme Binding Proteins GSBP1 and GSBP2, recently identified in *Spirostomum minus*, which contain many tandem repeats of the consensus sequence and have a very large size of up to 2000 kDa, more than 20 times larger than *S. cerevisiae* Sfi1 [5]. Of these, representative Sfi1 motifs and representative centrin were chosen for further analysis through AlphaFold and *in vitro* reconstitution.

These sequences show that the organization of *Spirostomum* Sfi1 is strikingly different from previously analyzed structures in *S. cerevisiae*. The sequences of the Sfi1-like proteins we identified in *Spirostomum* contain many consecutive 69-amino acid repeats with very few non-repeat interruptions (Table S3). Thus, the spacing of the expected centrin-binding sites is highly regular over a significant fraction of the protein’s length. In addition, the sequence signature of these repeats is different from the canonical Sfi1 repeat of *S. cerevisiae* and in other related eukaryotes. First, each *Spirostomum* 69-amino acid repeat includes two proline residues. In contrast, there are relatively few proline residues within the *S. cerevisiae* Sfi1 protein (see Supplementary Table S4). These prolines divide the 69-amino acid repeat into two sub-repeats, 36 and 33 amino acids in length. Second, a highly conserved tryptophan is observed in each repeat of Sfi1 we identify in *Spirostomum* (Supplementary Tables S3), but it appears less conserved in *the S. cerevisiae* Sfi1 repeats (Supplementary Table S4).

To examine the structural consequences of these sequence features, we used AlphaFold to predict the structure of a 15 repeat sequence from *Spirostomum ambiguum* (Supplementary Table S3) and a comparable number of repeats from *S. cerevisiae* Sfi1 (Fig. 5A, Supplementary Data 1 and 2, Supplementary Table S4) [38– 40]. The predicted AlphaFold structure of the *Spirostomum ambiguum* Sfi1 repeat includes significantly more kinks within the repeats compared to the yeast Sfi1 (Fig. 5A, Supplementary Data 2). These interruptions to an extended alpha-helix occur near the locations of helix-breaking amino acids, such as the proline residues noted above, as well as glycines. These differences suggest a much shorter persistence length for *Spirostomum ambiguum* Sfi1-like repeats than for *S. cerevisiae* Sfi1.

While not conclusive, these data suggest a model of contraction wherein calcium binding and unbinding from centrin might regulate the formation and stabilization of kinks not observed in yeast Sfi1, thereby stabilizing either a more elongated or a more compact state of the large Sfi1 protein. We attempted to validate this model by comparing the predicted AlphaFold structures in the presence and absence of calcium. While AlphaFold produces both elongated and compact Sfi1 structures with and without calcium (Supplementary Fig. S5), AlphaFold models are biased by both PDB training data and architectural assumptions toward finite oligomers with a defined number of chains and point-group symmetry, and are consequently less effective at modeling helical assemblies that require screw symmetry and periodic translation. Thus, we turned instead to reconstitution experiments.

### Myonemal protein complexes compact *in vitro* in response to calcium

While AlphaFold alone cannot provide conclusive evidence of *Spirostomum* centrin/Sfi1 contraction, its predictions led us to hypothesize that truncated *Spirostomum* centrin/Sfi1 complexes might compact in response to calcium, in turn driving myonemal contraction. Secondly, we hypothesized that meso-scale contraction of multi-filament complexes could be driven through multimerization or phase-separation interactions, again regulated by calcium, which could drive the long Sfi1 filaments into a more compact state. We sought to build an *in vitro* reconstitution of the myonemal centrin/Sfi1 complex to test both of these hypotheses *in vitro*.

To do so, we co-expressed a construct containing 3 tandem *Spirostomum* Sfi1 repeats (69 amino acids each), which we call Sfi1×3, and a second construct of *Spirostomum ambiguum* centrin, hereafter simply referred to as centrin, in *E. coli*. The Sfi1×3 construct was flanked by an SH3 domain and ligand, which improved purification yields, and both constructs contained a His-tag for purification (Figure 5A). These proteins expressed efficiently, and we were able to successfully purify the complex with NiNTA resin (Fig. S6). Next, we characterized *in vitro* activity in response to calcium of individual complexes by analytical ultracentrifugation (AUC) and larger multimers by fluorescence microscopy.

We characterized individual complexes by AUC in the presence of high salt (1 M NaCl) to prevent the formation of large complexes. We used two methods: the steadystate method, where diffusion and centrifugal forces balance each other to measure mass; and the dynamic velocity measurement, where hydrodynamic drag can be measured and differences can be attributed to changes in mass or changes in the complex shape. The mass measurement without calcium (Fig. S7A) yields a molecular weight of 121.8 ± 4.0 kDa and is most consistent with a complex of 4 centrins (20.9 kDa each) to one Sfi1×3 (36.6 kDa). While we cannot know if this purified complex reproduces the native stoichiometry, this measurement is consistent with 1-2 centrins bound per *Spirostomum* Sfi1 repeat, as predicted from the comparison with yeast Sfi1.

We then repeated this measurement with the addition of 10 mM CaCl_2_, and found a very similar primary mass component of 121.4 ±3.0 kDa (Supplementary Fig. S7B), suggesting that there is no change in mass between the two conditions that would be indicative of a change in centrin/Sfi1 binding. Additionally, we ran an experiment that subtracted these two conditions at the same radius and find a flat line with only small differences for the largest masses and not the primary component (Supplementary Fig. S7C). We also detected a secondary component with a mass of 830.5 ± 424.5 kDa without calcium and 901.7 ± 247.5 kDa with calcium. Additionally, the subtracted data show a slight increase in the heavier component after subtraction. However, since these larger complexes are only a small fraction of the total mass of the sample (15% and 13% without and with calcium, respectively), we focus primarily on the shape of the primary mass component in subsequent analyses.

We next used the AUC dynamic velocity measurement to assess hydrodynamic drag and to test for the compaction of individual complexes. From this experiment, we found that while the mass between the two samples remained nearly identical (indicating no change in stoichiometry), the sample with the addition of calcium tended to sediment significantly faster in the dynamic velocity measurement, indicating compaction with the addition of calcium (Fig. 5C). From the raw data, we were able to fit the sedimentation coefficients and average them between two repeats of sample preparation, with three AUC runs each, showing a statistically significant difference in the measured sedimentation coefficient (Fig. 5D). These data strongly support that individual centrin/Sfi1 complexes become smaller (more compact) in response to calcium, a behavior that we hypothesize could be one of the drivers of myoneme contraction.

Finally, we test for larger scale changes in inter-complex interactions that may cause an increase in bundle density during contraction, like those seen in Fig. 3. To do so, we fluorescently labeled the purified protein and imaged it with wide-field fluorescence microscopy. We found that there were significantly more very dense puncta under higher calcium conditions. This was in stark contrast to the low calcium conditions and the control, which showed little puncta formation and tended to be more uniform. To quantify these differences, we plotted histograms combining the pixel values to show a trend towards more extreme values as we increased the calcium concentration, indicating a transition from the protein being in solution to large scale aggregation (Fig. 5E), which begins at a physiologically-reasonable concentration of ~100 *µ*M. Some of the larger aggregates from each sample condition are shown in the representative images in (Fig. 5F). In total, the *in vitro* data lead us to propose a phase diagram (Fig. 5G) where increased calcium concentration drives both the compaction of individual centrin-binding Sfi1 repeats and self-association between repeats (likely driven by centrin-centrin associations), which further drives the compaction of the filament as a whole.

## DISCUSSION

The data presented herein, which span from the molecular to the organismal scale, lead us to propose a new multiscale model for how structural changes at the molecular level power the contraction of *Spirostomum* as a whole (Fig. 6). From its molecular triggering to its explanation of organismal force transduction, this proposed mechanism is notably distinct from better characterized cytoskeletal force generators, such as actomyosin contraction and microtubule sliding.

**FIG. 6:**
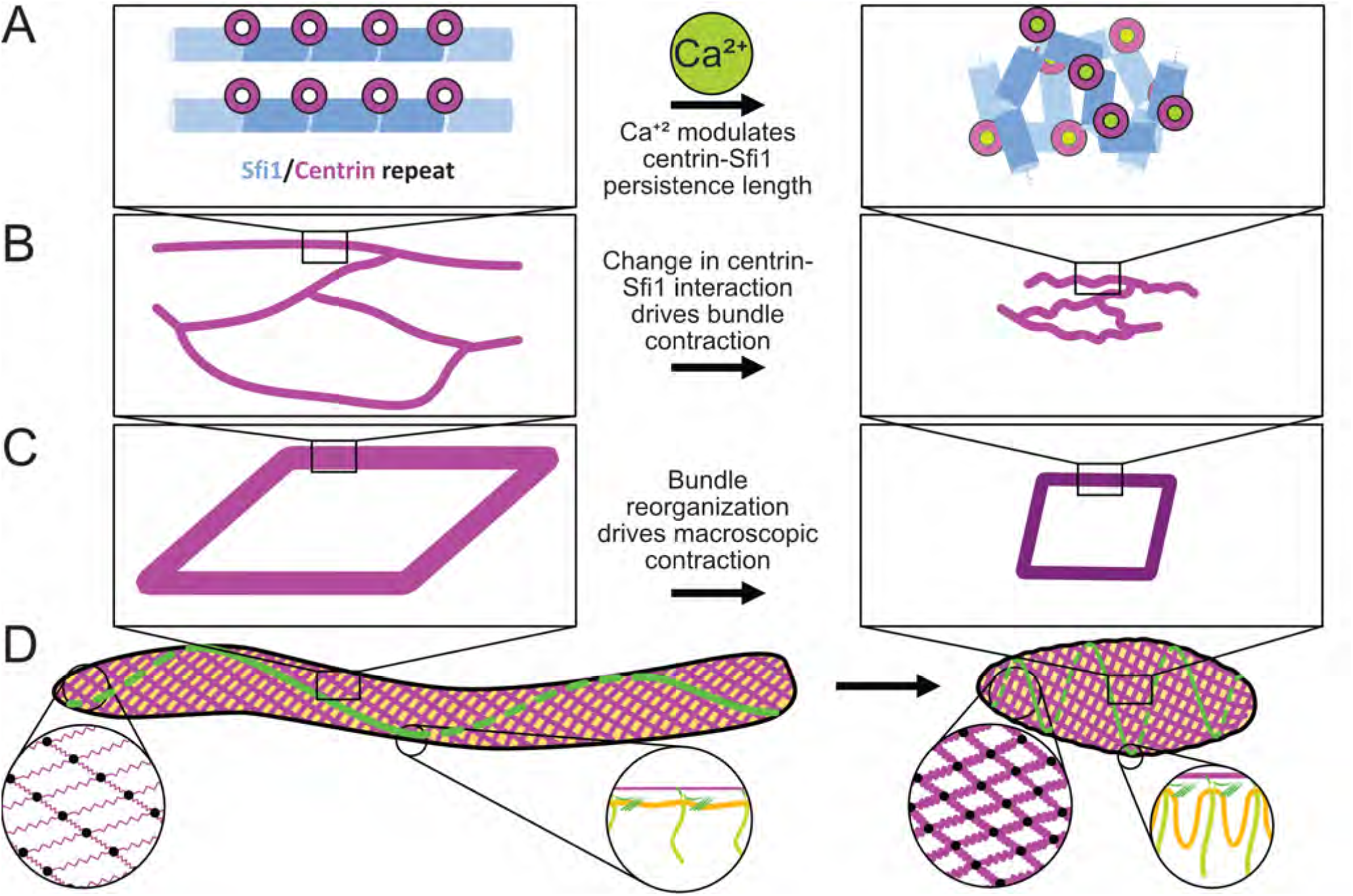
Proposed multiscale (nm-mm) hierarchical physical mechanism for Sfi1/centrin powered contraction in *Spirostomum*. **A** On the protein scale, we propose that calcium-signaled centrin modulates the stiffness of Sfi1 filaments. **B** At the ≈ µm scale, the change in persistence length of these filaments generates contractile force directed along the organism’s long axis. **C** At the mesoscale, myoneme filaments form the edges of the fishnet-like mesh, which contracts the cortex while preserving overall organismal structure. **D** In the organism as a whole, myonemes generate contraction. Meanwhile, membrane ridges, templated via anchorage at basal bodies to myonemes and cortical microtubule bundles, preserve overall surface area.

Reconstitution of centrin/Sfi1 proteins reveals initial insights into the molecular mechanism for myonemal contraction. We show that, in response to calcium, individual repeats of centrin/Sfi1 complexes compact while also forming higher order multimers. We propose that these two potential mechanisms may both help drive contraction: a compaction of individual units of the myoneme filaments leading to an end-to-end force, as well as the cross association of repeated units on the same or neighboring filaments, leading to shortening and an increase in density.

While higher resolution structural information is still needed to fully decipher the contraction of *Spirostomum* centrin and Sfi1, differences in the amino acid sequences of Sfi1 in *Spirostomum* compared to its *S. cerevisiae* counterparts may hint at how it achieves its distinct function. Modeling predicts that the helix-breaking residues allow kinks or bends in *Spirostomum* Sfi1 that disrupt the otherwise largely alpha-helical structure. Calcium regulation may take advantage of these regularly spaced helix breaking residues, driving *Spirostomum* Sfi1-centrin complexes into a more compact state (Fig. 6A). This compact state could be stabilized by specific protein/protein interfaces, or, alternatively, the Sfi1 filaments may act as an entropically driven spring whose stiffness is regulated by calcium (Supplementary Discussion 1). Note that Salisbury (who attributes his model to Szent-Györgyi) proposed a conformational change-driven mechanism for the force generated by Sfi1/centrin in yeast [22]. We do not rule out the contributions of such a model in *Spirostomum*, or of other specific rearrangements that may become clear once we have a fuller structural picture of this large protein complex. However, we do note that an entropic spring-like mechanism would seem to take advantage not only of the regular helix-breaking residues seen in *Spirostomum* Sfi1 homologs but also of the immense size that is their distinctive feature. This proposed mechanism has some similarity to the giant protein titin, which has also been proposed to act as an entropic spring to support passive relaxation and/or supplement contraction in the sarcomere [41]. However, generating organismal scale contraction by modulating filament persistence length would, to our knowledge, be both novel and unique in biology.

We can also now begin to understand how nanometer scale changes in myonemal proteins are transduced into bundles that contract at the micron scale (Fig. 6B). The mesh that we see in elongated myonemes has an approximately hexagonal network with irregular branching points. Furthermore, *Spirostomum* myoneme filaments appear to shorten perpendicularly to contraction. These rearrangements are markedly different from other large, repeatedly contracting cellular structures such as actomyosin filaments in muscles, which include highly uniform repeating bands that maintain a constant width as they contract [42]. Although irregular actomyosin networks can also contract [43], the sarcomeric architecture helps them to do so more robustly and efficiently [44]. An irregular fiber bundle formed from many filaments, whose persistence lengths are modulated by calcium, may allow *Spirostomum* to achieve efficient contraction with greater structural flexibility. Furthermore, the dense, multimerized interactions we see in response to calcium *in vitro* are consistent with the highly dense myoneme in contracted cells that we see by TEM.

Although the structure within the myoneme filaments appears somewhat random, more organization emerges on the organismal scale (Fig. 6C). The orientation of the myoneme mesh amplifies the force of each individual bundle segment, as they act in parallel when encircling the organism. Longitudinally, from head to tail, the filament bundles act in series, which allows for a small change in the size of each filament to add up to a large change in the overall length of the organism. Furthermore, the angled, fishnet mesh ensures uniform contraction without bulges or irregularities that might lead to stress failures of organelles or the organism as a whole. Together, these effects allow *Spirostomum* myonemes to generate large cortical forces that dramatically reshape the organism without destroying its internal structures, and to do so in a manner strikingly distinct from other, more characterized mechanisms of biological contraction (Fig. 6D).

When myonemes contract, the immense loads accompanying their rearrangements must be effectively dissipated and productively distributed to other cellular structures to alter the shape of the organism while preserving its contents [45]. The regular structures we observe near the cell cortex in contracted cells, namely, bent microtubule bundles and membrane ridges, may help ensure that the force of contraction is directed into structures that are strong and flexible enough to withstand it. The formation of these structures is not yet fully clear, but they may be patterned via mechanical coupling between the myonemes and the plasma membrane. With the basal bodies of the cilia connected together by microtubules acting as anchors, the membrane can buckle into uniformly distributed ruffles, much like a shower curtain being held by hangers, with the force generated by the myoneme. The association we observe between myonemes, cortical microtubule bundles, and the basal bodies of the cilia may provide this coupling.

We have proposed here a mechanical framework that begins to explain the rapid contraction of *Spirostomum* and its molecular mechanism. However, important questions remain. First, the mechanism of elongation following *Spirostomum* contraction is yet completely uncharacterized. Although we explore the idea that microtubule bundles and the cell membrane may store energy from contraction, we estimate that the relevant energies are orders of magnitude apart in scale, leaving them insufficient to explain elongation. Thus, we think it is likely that as yet undescribed mechanisms allow this organism to elongate. One possible mechanism for elongation is the force generated by the cilia, which contributes to the neck extension in *Lacrymaria olor* [46]. Another possibility is that the active re-sequestration of calcium allows the myonemes themselves to produce an elongation force.

Regarding contraction, while centrin and Sfi1 are very likely key force generating myonemal components, further assays would help to conclusively demonstrate the roles of filament compaction and aggregation in force generation, or whether additional components are required to entirely explain their force generation. If they are sufficient, it will still be important to identify the specific interactions that drive both contraction and elongation within the Sfi1/centrin interaction. The exact details of the aggregation/multimerization, including whether liquid- or gel-like, also merit additional investigation. Centrins from various organisms have previously been shown to undergo calcium-induced polymerization *in vitro* [47, 48], *and this behavior has been proposed to be liquid-liquid phase separation, which may be important in malaria infection [49]. Similarly, it is important to note that there are multiple centrins and Sfi1 motifs within Spirostomum*, and the ones we have characterized provide only the first insights into their behavior. New molecular and cell biological tools will be needed to fully elucidate how this diverse biochemistry might contribute to *Spirostomum* structure and behavior, but the possibilities are intriguing. For instance, both the three-branch nodes and the centrin-containing filaments that we observe near basal bodies by TEM provide important clues that additional interactions and/or biochemical components are critical for myoneme organization. Some of the many distinct *Spirostomum* centrins might localize to these and could, in turn, recognize specific regions of Sfi1.

We have demonstrated that myoneme filaments are key in myoneme force generation and mechanistically diverge from the motor-driven systems that constitute a prevailing mechanism of contractile force generation in biology. This form of contraction offers new perspectives for biological models and engineering designs on both macro and microscopic scales.

## MATERIALS AND METHODS

### Cell culture

*Spirostomum* media was made by boiling 1 L of spring water (Carolina Biological Item #132450) with 10 wheat seeds (Carolina Biological Item #132425) and a few leaves of Timothy hay (Carolina Biological Item #132385)[50]. We obtained initial *Spirostomum ambiguum* cultures from Carolina Biological (Item #131590), which were confirmed by scientists there to be *Spirostomum ambiguum*. New cultures were created by inoculating the media with cells from a mature culture. Mature cultures that had at least two weeks to grow were used for all experiments, since after this time the *Spirostomum* organisms we observed were longer and more consistent in appearance than those from cultures that were recently inoculated.

### Fixation and immunoflorescence microscopy

Cells were fixed using two methods: slow PHEM-formaldehyde and fast EGTA-Methanol fixation. No differences were found in the localization of the two methods for fixation; however, the methanol fixed cells were somewhat more robust to handling after fixation.

#### PHEM-formaldehyde fixation

Cells were transferred from cultures into a 96-well plate and washed with spring water twice. To obtain elongated samples, cells were incubated in 200 µL of 0.5X PHEM buffer (1X PHEM buffer: 60 mM PIPES, 25 mM HEPES, 10 mM EGTA, and 2 mM MgCl_2_, pH 6.9 [51]) at 4°C for 6 hours and then fixed for 1 hour with 100 µL of 1X PHEM 2% PFA at 4°C. To produce contracted samples, cells were rinsed in 100 µL of 1X PHEM buffer at room temperature (RT) for 15 seconds and fixed with 100 µL of 1X PHEM 2% PFA for 15 minutes. The cells not treated with EGTA (i.e. PHEM buffer) consistently contracted upon contact with the formaldehyde solution, likely due to either chemical or mechanical stimulation. Cells exposed to EGTA under the fixation conditions exhibited no visible contraction as confirmed by observing them as they were exposed to fixatives under dissection microscopes. Fixed cells were then treated with 300 µL of 2% PFA 1X PHEM for 60 minutes and washed three times with 300 µL of permeabilization solution (1X PBS, 3% BSA, and detergent, which was either 1% saponin or 0.1% Tween) for 10 minutes each. After fixation, cells were stained for immunofluorescence (see below). Note that some mucus-like debris is generally present in fixed samples observed on the surface of the cells.

#### Formaldehyde-methanol fixation

The second method for fixation was a fast incubation in EGTA in spring water, followed by iced methanol fixation. Methanol fixed samples were more stable to mechanical perturbation, degraded less by permeabilization, and were less likely to be damaged from handling and repeated washings.

Cells were transferred from the culture into a 1.5 mL tube and rinsed 3 times with spring water. The elongated cells were prepared by exposing the cells to 25 mM EGTA in spring water for about 30 seconds, until the cells no longer contracted in response to mechanical tapping of the tube. Contracted samples skipped this step and went straight to fixation. Cells were then dropped into −80 °C methanol with 2% formaldehyde (prepared from 16% formaldehyde/H_2_O). Without EGTA, the cells would contract, likely due to either chemical stimuli or the force of the droplet hitting the fixative. The samples were kept at −80 °C for 1 hour, −20 °C for 1 hour, and finally left in 4 °C overnight. The samples were rinsed in a 1:1 solution of methanol:PBS. followed by rinsing 3 times in PBS. Cells were then transferred to a glass bottom dish (MatTek P35GC-0-14-C) for staining.

#### Immunofluorescence staining and sample mounting

To visualize cytoskeletal components by immunofluorescence, the following primary antibodies were used: myoneme: 20H5 mouse anti-Centrin (Millipore Sigma, Cat. #04-1624) (previously used to identify centrin in ciliates [52]) or Sfi1 antibody [19]; microtubules: TAP952 mouse anti-alpha tubulin (Millipore Sigma, Cat. #MABS277). Cells were incubated in 200 µL primary antibody solution diluted to 10 µL/mL in permeabilization solution, incubated overnight at room temperature (RT), and washed three times with 300 µL of permeabilization solution for 10 minutes each. Secondary antibodies (ThermoFisher, Cat. #R37115, A11008, and A21244) were diluted 1:1000 in permeabilization solution to prepare the secondary solution. Cells were incubated in 200 µL of secondary solution for three hours at RT and washed three times with 300 µL of 1X PBS 3% BSA for 10 minutes each before imaging. To stain the cell membrane, cells were incubated in 200 µL 1:1000 Cell-Mask Orange Plasma Membrane Stain (ThermoFisher, Cat #C10045) in PBS for 1 hour and rinsed 3 times in PBS before imaging.

For microscopy, stained cells were either transferred onto a glass cover-slip (Neuvitro, Cat. #GG-22-1.5-PLL) with 100 µm spacers (Cospheric, CPMS-0.96 106-125 µm) and a glass slide (VWR, Cat. #16004-430) and sealed with nail polish (Fisher Scientific, Cat. #72180) to prevent evaporation, or samples were placed in a glass bottom dish (Maktek P35G-1.5-14-C) with a cover-slip placed over the well to prevent evaporation.

The 20H5 antibody was validated by in gel western blot against both recombinant and native *Spirostomum*. 1 mL of dense *Spirostomum* culture was washed three times in spring water before being mixed with 25 mL of 4x Laemmli Sample Buffer (Biorad #1610747), 10 mL of Protease inhibitor cocktail (ThermoFisher #PI87786) and brought to a final volume of 100 mL then heated to 95 °C for 15 minutes. Then, it was run on a 4%-20% gradient gel (Biorad #4561093EDU); various quantities of protein were loaded, as well as the purified construct below (see Supplementary Fig. S2); the gel was run for 15 minutes at 50 V and then at 120 V until the dyes were near the bottom of the gel. The gel was then washed in DI water for 30 minutes. The gel was then fixed in 50% methanol and 5% acetic acid overnight. The gel was then blocked with 3% BSA, 25 mM Tris-HCl, pH 7.5, for 1 hour with shaking. It was rinsed with the same buffer, then incubated with 1:10000 20H5 in 3% BSA, 25 mM Tris-HCl, pH 7.5, shaken for 1 hour, and then placed overnight in 4 °C. The gel was rinsed and then placed in 1:10000 Licor IRDye 680RD Goat anti-mouse IgG for 2 hours, shaking at room temperature. It was finally rinsed three times in TBST and imaged on a LICOR odyssey clx.

#### Fluorescence imaging

Cells were imaged by immunofluorescence microscopy on the following microscopes using Z-stacking and Mosaic stitching: Zeiss LSM 780 Elyra PS1 Super-resolution equipped with Zeiss Plan-Apochromat 20x/0.8 M27 objective, excited with 555 and 594 nm lasers, and with filters 60-800, 599-734, MBS 458/514/594; Nikon Ti-E Dragonfly Spinning Disk Confocal equipped with Nikon Plan Fluor 20x/0.75 Mlmm, Nikon Plan Apo 100x/1.45, and Nikon Apo TIRF 60x/1.49 objectives, Andor Zyla sCMOS and Andor iXon EMCCD cameras, excited with 488, 561, and 637 nm lasers, and with Chroma filters ET525/50m, ET600/50m, ET700/75m, ZT405/488/561/640rpc; Nikon Ti2 Eclipse AX/AX R NSPARC equipped with Nikon Plan Apo λD 60x/1.42 objective, excited with 488, 561, and 640 nm lasers, and with filters 502–546 nm, 570–616 nm, 666–732 nm, 405/488/561/640.

### Sfi1 antibody generation

Custom anti-Sfi1 peptide antibodies (Bethyl Laboratories, Montgomery, TX) were generated for selected peptides identified in *Stentor*. Rabbit preimmune bleeds were first screened to avoid using rabbits that may have been previously exposed to ciliated parasites and already produced antibodies that react with *Stentor* proteins in immunofluorescence. Sfi1 sequence reads were obtained from *Spirostomum* and aligned against *Stentor* Sfi1 to identify homology. The following *Stentor* Sfi1 peptide sequence was identified as having the highest homology to *Spirostomum ambiguum* Sfi1: RTEKLRNAL-NRVPR (Supplementary Table S1). The antibody raised against this peptide was also identified as being reactive to *Spirostomum ambiguum* Sfi1 by colocalization in immunofluorescence with centrin (Supplementary Fig. S3). Validation of the antibody in Stentor was performed by Yan et al. [19].

### Transmission electron microscopy

TEM methods have been adapted from previously published methods [53, 54], as described below.

#### Fixation and Embedding

Contracted samples were fixed in a fixation buffer of 2.5% glutaraldehyde (EMS Cat # 16220) and either PBS (ThermoFisher CAT # 70011044) or cacodylate buffer (100 mM, pH 7.4) [53]. Samples were first washed three times in spring water. Elongated cells were prepared by exposing the cells to 25 mM EGTA in spring water for about 30 seconds, until the cells no longer contracted in response to mechanical tapping of the tube. Contracted and elongated cells were then taken in a small droplet and dropped into 2 mL of fixation solution and incubated for 2 hours on ice. After fixation, samples were rinsed in glutaraldehyde-free buffer and then subjected to an ethanol series (50%, 75%, 90%, 100% dilution by volume of ethanol with buffer for 1 hour each) to slowly dehydrate the samples. Samples were left in 100% ethanol overnight before a final rinse in 100% ethanol and then a resin series, where they were left in 50% LR white (EMS CAT # 14380) overnight, followed by 1 hour each of 75%, 90%, and then 100% (resin diluted with ethanol); they were then placed in gelatin capsules (EMS CAT # 70110) and baked at 70°C overnight [53].

#### Microtomy

The samples were microtomed on the Leica UC7 with a glass or diamond knife to a thickness between 70 and 120 nm. The sections were then placed on uncoated 200 mesh nickel grids (EMS CAT# 200-Ni or H200-Ni). We found that the immunolabeled samples prepared using uncoated grids were more fragile and more likely to form holes under the electron beam (as seen in Fig. 3); however, the labeling was far superior, likely due to the two exposed surfaces of the section during staining [55].

#### Immunogold labeling

Immunogold labeling was performed to label centrin in samples according to EMS protocols [56]. The incubation solution was prepared with 20 mM PBS (Fisher Scientific #BP399500), 0.1% Aurion BSA-C (SKU: 900.099), and 15 mM sodium azide (CAS 26628-22-8). Following fixation, embedding, and microtomy, as described above, the sectioned samples were first washed with 50 mM Glycine (CAS 56-40-6) in PBS for 15 minutes to inactivate any remaining glutaraldehyde. They were then incubated in goat blocking solution (Aurion SKU: 905.002) for 15 minutes and rinsed twice in the incubation solution. The grids were then left in 5 µg /ml 20H5 mouse anti-centrin antibody (Millipore Sigma, Cat. #04-1624) or Sfi1 antibody (see above) overnight at 4°C. Secondary only controls were performed in the same way, with the exception of the primary antibody addition to the incubation solution (Supplementary Fig. S8). The next day, the sections were rinsed five times in the incubation solution and then incubated in 1:100 10 nm gold-conjugated secondary antibodies (abcam cat #: ab39619 or Invitrogen Ref: A31566) in the incubation solution for 2 hours. The samples were then rinsed five times in the incubation solution, twice in PBS, and post-fixed in 2.5% glutaraldehyde for 15 minutes before being rinsed three times in distilled water. The samples were then stained with 4% uranyl acetate (CAS 541-09-3) for 15 minutes and rinsed three times in distilled water. The samples were then imaged on the Hitachi HT7800 after drying.

### Identification and expression of myoneme proteins

Detergent-extracted, taxol-stabilized cytoskeletons were prepared from *Spirostomum* cultures, and their proteins were separated using SDS-PAGE. Bands within the expected centrin size range (20-30 kDa) were excised and analyzed by mass spectrometry. No protein sequence data were available; RNA-seq data for this species were retrieved from the NCBI SRA database (SRR12125180) and processed with the Trinity and rnaSPAdes tools on the Galaxy platform [57]. The assembled transcriptomes were translated using Virtual Ribosome 2.0 [58] to derive the *Spirostomum ambiguum* proteome. Using this derived proteome, mass spectrometry of tryptic peptides from the excised bands identified several dozen proteins. A BLAST search was performed using the *Paramecium tetraurelia* infraciliary lattice protein ICLa (GenBank: AAC47156.1) as a query because ICL proteins are known to be centrins that bind to Sfi1-like scaffold proteins, forming a Ca2+-driven contractile assembly within the cortex of *Paramecium* [59]. Two top-scoring hits were identified in the mass spectrometry data. These ICL1a homologs in *Spirostomum* were nearly identical to those found in *Stentor* and *Blepharisma*, two other heterotrich ciliates that exhibit myoneme-dependent contractile behavior. One of these *Spirostomum* proteins was selected for further study, and a synthetic gene encoding a Histagged version was optimized for expression in *E. coli* (ATUM). This gene was cloned into the pJ414 plasmid (ampR), then transformed and expressed in *E. coli* BL21 (DE3). A representative 15 repeat section of *Spirostomum* Sfi1 was selected for analysis using AlphaFold in conjunction with the selected centrin sequence.

To identify candidate Sfi1-like proteins in *Spirostomum ambiguum*, BLAST searches were performed using one of the *Paramecium tetraurelia* giant centrin-binding proteins (GenBank: CAI39100.2), which contains multiple Sfi1 repeats as the query. Among the high-scoring hits were several proteins with a variable number of tandem conserved 69 amino-acid repeats. Based on these sequences, a synthetic gene was constructed encoding three tandem, non-identical 69-amino-acid repeats (Sfi1×3). Genomic sequencing of *Spirostomum minus* by other researchers indicated that these repeats are likely derived from fragments of a much larger gene [5]. To promote the self-assembly of this protein into a longer centrin-binding scaffold, a His-tagged N-terminal SH3 domain and a C-terminal SH3 ligand sequence were added. This construct was cloned into a pJ411 (kanR) plasmid and transformed into *E. coli* BL21 (DE3). However, expressing this protein alone resulted in low yields. Therefore, *E. coli* cells were transformed with both the centrin and Sfi1 repeat plasmids, and co-transformants were selected on LB agar plates containing ampicillin and kanamycin. Co-expression with centrin improved the stability of the Sfi1 repeat protein, enabling substantial production of both proteins in the same cell. Additionally, a control construct was created with the same SH3 domain and ligand sequence, using a rigid spectrin linker and expressed in pET-21a(+) (Novagen).

Cell lysates from cells co-expressing centrin and Sfi1×3 proteins were treated with B-PER Plus bacterial cell lysate solution (Thermo Fisher), and the proteins in the post-lyse pellet were solubilized using a 1 M NaCl extraction buffer. The proteins were then purified using HisPur Ni-NTA or cobalt spin columns (Thermo Fisher) with buffers containing 1 M NaCl. Pooled eluted fractions were concentrated and buffer-exchanged on PD10 desalting columns (Cytiva) to remove imidazole and phosphate and to add the calcium chelator EGTA. An SDS page gel showing samples from each stage can be seen in Supplementary Fig. S6.

For labeling, the proteins were exchanged into 1 M NaCl, 20 mM sodium phosphate, pH 7.4. Equimolar DyLight 650 NHS Ester (Thermofisher 62265) was added and allowed to react overnight at 4 °C. The resulting protein was then exchanged into 1 M NaCl, 25 mM Tris-HCl, pH 7.5, and 1 mM EGTA for storage, which also removed any excess dye (Fisher Sci. PI89883). For imaging, the protein was exchanged into 25 mM tris, pH 7.5, 5 µmol L^−1^ EGTA (Fisher Sci. PI89883).

96 well glass bottom plates were prepared for imaging by rinsing three times with 100% ethanol and then with milliQ water. This was then dried with filtered compressed air. The wells were then blocked with 3% BSA and 25 mM Tris-HCl,5 µmol L^−1^ EGTA for 30 minutes while rocking. This was rinsed with MilliQ water. The wells then had 1 µL of protein added to the bottom (Cen/Sfi1 1: 0.5 mg/mL, Cen/Sfi1 2: 0.4 mg/mL, Sh3 Control: 0.15 mg/mL, Measured by nanodrop before labeling). 1 µL of the two Cen/Sfi1 constructs and 3 µL of the control were used to bring the concentration halfway between the other two for the control sample. Then, 100 µL of 1% BSA and 25 mM Tris-HCl, with the addition of calcium chloride and magnesium chloride, was added for each of these conditions: 0 mM CaCl2, 10 mM MgCl2, and 5 µM EGTA; 0.1 mM CaCl2 and 9.9 mM MgCl2; 1 mM CaCl2, 9 mM MgCl2; 10 mM CaCl2, 0 mM MgCl2

Preliminary experiments showed this to be the relevant sensitive range of concentrations of calcium and magnesium ions that were added to control for the total concentration of divalent cations in the buffers. The solutions were allowed to shake for 15 minutes before imaging. Imaging was then performed using EPI-fluorescence on the Nikon TI-Eclipse 60x (See fluorescence imaging above for details). Images were taken in a 10×10 mosaic pattern. 2 independent preparations of protein were used, with two sample preparations of each.

### Analytical ultra-centrifugation of constructs

Two independent preparations of the centrin/Sfi1 constructs were used to perform sedimentation velocity experiments, with each experiment being repeated three times. Samples and buffer were loaded into 2-sector centerpieces that were 1.2 cm in height and were run in a ProteomeLab XLA analytical ultracentrifuge (Beckman Coulter) using an AN60-Ti rotor. The cells were spun at 40,000 rpm, and absorbance data at 280 nm were collected over 400 scans per cell.

A sedimentation equilibrium experiment was also performed to measure the mass under both conditions. We used a 6-sector centerpiece that is 1.2 cm in height and loaded the sample and buffer into the first four sectors. The final sectors contained our EGTA protein sample in the buffer sector and our CaCl2 protein sample in the sample sector. The samples were allowed to come to equilibrium at 7,000, 8,500, and 10,000 rpm, during which we collected scans at 280 nm in intensity mode. Buffer Conditions: 25 mM HEPES, pH 7.5, 1 M NaCl, 10 mM Calcium or 10 mM EGTA. Final Protein concentration measured: 0.15 mg/mL.

Sedimentation velocity data were analyzed using Sedfit version 17.0 with the Continuous c(s) and Continuous c(s,ff0) models [60, 61]. Buffer density and viscosity were calculated using SEDNTERP [62, 63]. For EGTA, we used the values for EDTA. Data visualization was aided by GUSSI [64].

Sedimentation equilibrium data were analyzed with Sedphat version 15.2 (SEDPHAT – a platform for global ITC analysis and global multi-method analysis of molecular interactions) using the Species Analysis with Mass Conservation Constraints model. We fit the data, allowing for two species.

### Quantification of light and electron microscopy images

Quantitative analysis was carried out primarily using ImageJ Fiji [65]. The lengths and angles obtained from immunofluorescence and TEM were measured using the measurement tools in ImageJ. TEM quantification was performed on the centrin-labeled samples in ImageJ by measuring the bundle width across multiple images and organisms, with 3 embedding repeats and over 10 labeling repeats. The tag density was measured by manually counting the gold nanoparticle labels on the myoneme and measuring the area over the myoneme in ImageJ using the polygon tool, where each point represents the average density over a single image. Cell lengths were measured by approximating a spline through the central axis of z-projected images labeled with CellMask Orange in ImageJ. Fluorescence imaging of constructs was analyzed in python using the numpy package [66]. The plots were created using the Python packages matplotlib and seaborn [67, 68]. For each immunofluorescence quantification, measurements from N individual cells (reported for each measurement) were averaged. These cell averages were then averaged to calculate the reported mean and standard error of the mean (SEM) over N samples, as reported in the figures and text.

### Skeletonization analysis

Skeletonization was carried out on the TEM images by first segmenting the structures using Ilastik [69]. The images were first reduced in resolution by 4x in each axis. Segmentation used an AI model to separate myoneme structures from the background by hand training the model and making corrections that were fed back into the model. Validation was carried out by overlaying the original images with the segmented and skeletonized images to ensure quality. Note that some of the filaments appear to change thickness, likely due to the filaments coming in and out of the plane of the section. This limits the precision of the skeletonization. The segmentation was then skeletonized in ImageJ Fiji [65]. The resulting skeletonized images were processed in a Jupyter notebook with OpenCV [70, 71]. Skeletonized images were first convolved with the following matrix:

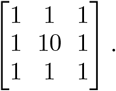

The resulting image could then be classified by the number of neighbors each pixel had. Pixels were classified by value as follows: 11 for endpoints, 12 for branches, and 13 to 18 for intersection points. Multiple nearby intersections could then be grouped into a single intersection. To extract edges, a branch traversal algorithm was used to find the branches that connected the intersections. From these data, we were able to calculate junction angles by counting out 5 pixels from each intersection along a branch, culling any smaller branches from the measurement, and calculating the angle between all of these points from the intersection. The lengths were also calculated for each branch using the Pythagorean theorem to determine the end-to-end distance between intersections, as well as the total length of the branch by counting pixels along the branch, adding 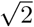 for a diagonal step or 1 for an orthogonal step. The lengths were then scaled by the pixel size in nanometers per pixel.

Following skeletonization, branch lengths were fitted to a worm-like chain model to estimate the persistence length of the branches:

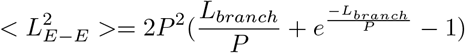

where L_*E−E*_ is the end-to-end distance, L_*branch*_ is the total length, and P is the persistence length [32]

### Coarse-grained mesh model

The mesh model was used to simulate three-dimensional changes in organismal shape in response to contraction. The model comprises N points **p**_*i*_ and an associated energy function 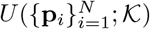, where 𝒦 represents a set of parameters. To mimic the contraction of the myoneme, which arises from a calcium wave that causes the centrin-based edges of the myoneme to shorten, we assume that calcium activation causes a change in the parameters 𝒦^elongated^ → 𝒦^contracted^. The dynamic aspects of the contraction used in a simplified continuum were studied in Floyd et al. [26]. Here, we study equilibrated structures following contraction and assume that for a given 𝒦, the energy is minimized with respect to the degrees of freedom **p**_*i*_.

Points 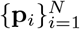 make up a quadrilateral spring mesh that encloses an elongated, cigar-like shape. The mesh is generated using the Catmull-Clark algorithm to sub-divide an initial rectangular prism into an iteratively finer surface [72]. The initial prism consists of n_*L*_ = 14 unit cubes stacked in the *z* direction (i.e., along the long axis of the cells) and centered on the origin. Points are then added to create progressively smaller quadrilateral faces. We run this process for three iterations (producing N = 3, 714 points), and following this initial mesh generation, we scale each coordinate so that the radius of the cylinder is 50 µm. In addition, the entire structure is given a z-dependent rotation so that each point is rotated in the xy plane by an angle:

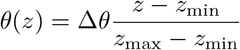

θ(*z*) linearly increases from 0 to Δ*θ* as *z* increases from the bottom to the top of the structure. We finally apply compression in the *z* direction to set the total length of the cylinder, ensuring that the angles of the unit cell in the fishnet mesh agree with the measurements on elongated myonemes. The result of this process is the latitudinal mesh (Fig. 2B), in which the edges lie along the lines of latitude of the cylinder. To produce the fishnet mesh, we move the latitudinal edges so that they connect to points between two adjacent layers and are offset by one relative to the original longitudinal edges.

Quadrilateral faces define the edges between the points **p**_*i*_ that are converted to springs in the energy function. We use a Hookean energy function for the spring 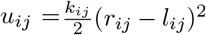, with *r*_*ij*_ = ||**p**_*i*_ − **p**_*j*_||. Before contraction, the rest lengths *l_ij_* are set as the initial distance between **p**_*i*_ and **p**_*j*_ in the mesh, ensuring that the system is in equilibrium. The new rest lengths are set as a factor γ times the original rest length: *l*_*ij*_ ← *γl*_*ij*_. The meshes can be made anisotropic by distinguishing between the two sets of springs (latitudinal and longitudinal in the latitudinal mesh, and right-handed and left-handed in the fishnet mesh) and assigning different shrinking factors for each direction. To assign the stiffness *k*_*ij*_ of each spring, we use the formula *k* = *EA*/*l*, where *EA* is the product of the Young’s modulus of the spring material and the cross-sectional area, and l is the length of the spring [73]. We assume a fixed value of *EA* for all springs of a given type and divide by the spring’s elongated rest length to obtain *k*_*ij*_. The total spring energy is

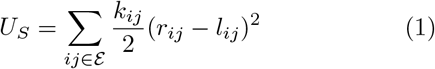

where ℰ is the set of edges in the mesh.

Because volume *V* is likely to be conserved during contraction, we constrain volume in the energy function *U*. We omit the conservation of the surface area here because the area can become ruffled at finer length scales than we resolve in the model (Fig. 1A, C). To calculate the enclosed volume, we create a surface triangulation 𝒯 by dividing each quadrilateral face of the latitudinal mesh into two oriented triangles. We can pick any point **o** in space (for convenience, we choose the origin) and sum the volumes of the tetrahedra that are defined for each face from the four points **o, p**_*i*_, **p**_*j*_, **p**_*k*_ [31]. The volume of the tetrahedron is

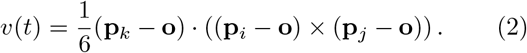

This will have a sign depending on the ordering of *i, j, k*, so we ensure that the orientation of each triangle *t* is the same for all *t* ∈ 𝒯. With this, the total volume is

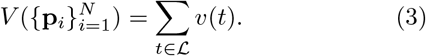

To incorporate *V* into the energy function *U*, we assign a rest volume *V*_0_ equal to the value in the elongated mesh. We then include

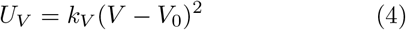

in the energy during minimization.

It is possible that there is resistance to torsion within the *Spirostomum* cortex. For instance, this could be attributed to the cortical microtubule bundles, which undergo additional bending if there is a twist between vertically adjacent layers of the mesh. To recapitulate this behavior, we treat adjacent layers as torsional springs. These springs penalize the relative twist Δψ_*l*_ and vertical separation Δ*z*_*l*_ between adjacent layers *l* and *l* + 1. The energy for the *l*^th^ layer is

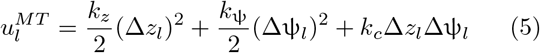

where *k*_*c*_ allows for coupling between stretching and twisting [73, 74]. The total energy of the microtubule is then a sum over the layers ℒ.

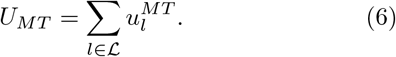

The stiffness parameters *k*_ψ_, *k*_*c*_, and *k*_*z*_ also scale inversely with the initial separation Δ*z*^0^ between layers. However, because Δ*z*^0^ = 8.2 µm is approximately constant throughout the elongated mesh, we directly parameterize the stiffness values. In this paper, we set *k*_*c*_ = *k*_*z*_ = 0 and focus on the effect of *k*_ψ_ in resisting the untwisting of adjacent layers in the latitudinal mesh; however, future work could explore the dependence of this coupling term on the dynamics of contraction and twisting of *Spirostomum*.

The total energy function is

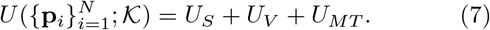

To mimic the effect of calcium activation, we set the new spring rest lengths and numerically minimized the energy function with respect to the points **p**_*i*_. To exclude edge effects, only the middle 95% of layers contributes to *U*_*MT*_ and has their rest lengths changed during contraction. Additionally, only the edges of these middle layers are rotated when converting the latitudinal mesh into the fishnet mesh.

To parameterize the model, the size of the organism was estimated to be approximately 100 *×* 1, 000 µ*m* in width and length. The initial twist of the mesh was also estimated from experimental images. Volume stiffness *k*_*V*_ was set high enough to ensure essentially perfect volume conservation. Bottom-up parameterization in terms of protein-level information is challenging because of the highly coarse-grained nature of the model. As a result, default values for the remaining parameters were obtained by searching the parameter space for minimized structures that reasonably matched experimental images. We then explored nearby regions of these default values to study the effect of these parameters on the observed behavior (Fig. 2). The default parameters are reported in Supplementary Table S2.

To quantify changes in concavity, we projected the 3D structure onto the longitudinal plane through the center of the structure and computed the projected area *A*_*p*_ in this plane. We then found the area of the projected shape’s convex hull *A*_*h*_ and computed the inflection ratio *R*_*ph*_ = *A*_*p*_/*A*_*h*_, which will be less than 1 for a non-convex structure.

### AlphaFold predictions of Sfi1 and centrin structures

*S. cerevisiae* sequences were obtained from the Saccharomyces genome database, SGD:S000003926 [75]. Predictions for *Spirostomum* and *S. cerevisiae* were generated using ColabFold v1.5.5: AlphaFold2 using the MM-seqs2 bot individually and in combination, as well as the AlphaFold3 server hosted at alphafoldserver.com [38–40, 76]. The AlphaFold2 initial models were relaxed using the built-in Amber software. The predicted structures were analyzed and visualized using the UCSF Chimera software [77].

## ACKNOWLEDGMENTS

The authors thank Fred Chang, Jane Maienschein, Scott Coyle, and members of the Bhamla, Elting, Honts, Marshall, and Dinner Groups for their feedback and helpful discussions; Aaron Bell for his help and training in electron microscopy and for his advice on the analysis of the TEM data; and Christina Hueschen and Alexander Kemper for their critical reading of the manuscript.

CF acknowledges support from the University of Chicago through a Chicago Center for Theoretical Chemistry Fellowship. This work was supported by the National Science Foundation under award numbers 1935260 and 2313722 (to MWE), 2313724, 1935262, and 1817334 (to SB), 2313727 (to JH), and 2313725 (to AD), and by NIGMS of the National Institutes of Health under award numbers R35GM130327 (to WFM) and R35GM142588 (to SB).

The authors acknowledge technical and instrumental support from the NC State Cellular and Molecular Imaging Facility (CMIF), the NC State Molecular Education, Technology and Research Innovation Center (METRIC), and the NC State Analytical Instrumentation Facility (AIF), all of which are supported by the State of North Carolina. AIF is also supported by the National Science Foundation (award number ECCS-2025064) and is a member of the North Carolina Research Triangle Nanotechnology Network (RTNN), a site in the National Nanotechnology Coordinated Infrastructure (NNCI). The authors also acknowledge instrument support from the Georgia Tech microscopy core. The authors acknowledge the University of Chicago’s Research Computing Center and the North Carolina State University High Performance Computing Services Core Facility (RRID:SCR_022168) for computing resources. Molecular graphics and analysis were performed with UCSF Chimera, developed by the Resource for Biocomputing, Visualization, and Informatics at the University of California, San Francisco, with the support of NIH P41-GM103311

## DATA AVAILABILITY

The assembled transcriptome will be made available before publication and is available upon request. All other data supporting the findings of this study are available within the paper and its Supplementary Information.

