## Supplemental Information for "A centrin–Sfi1 myoneme fishnet powers ultrafast calcium-triggered contraction in the giant ciliate *Spirostomum ambiguum*"

**SUPPLEMENTAL FIGURES**

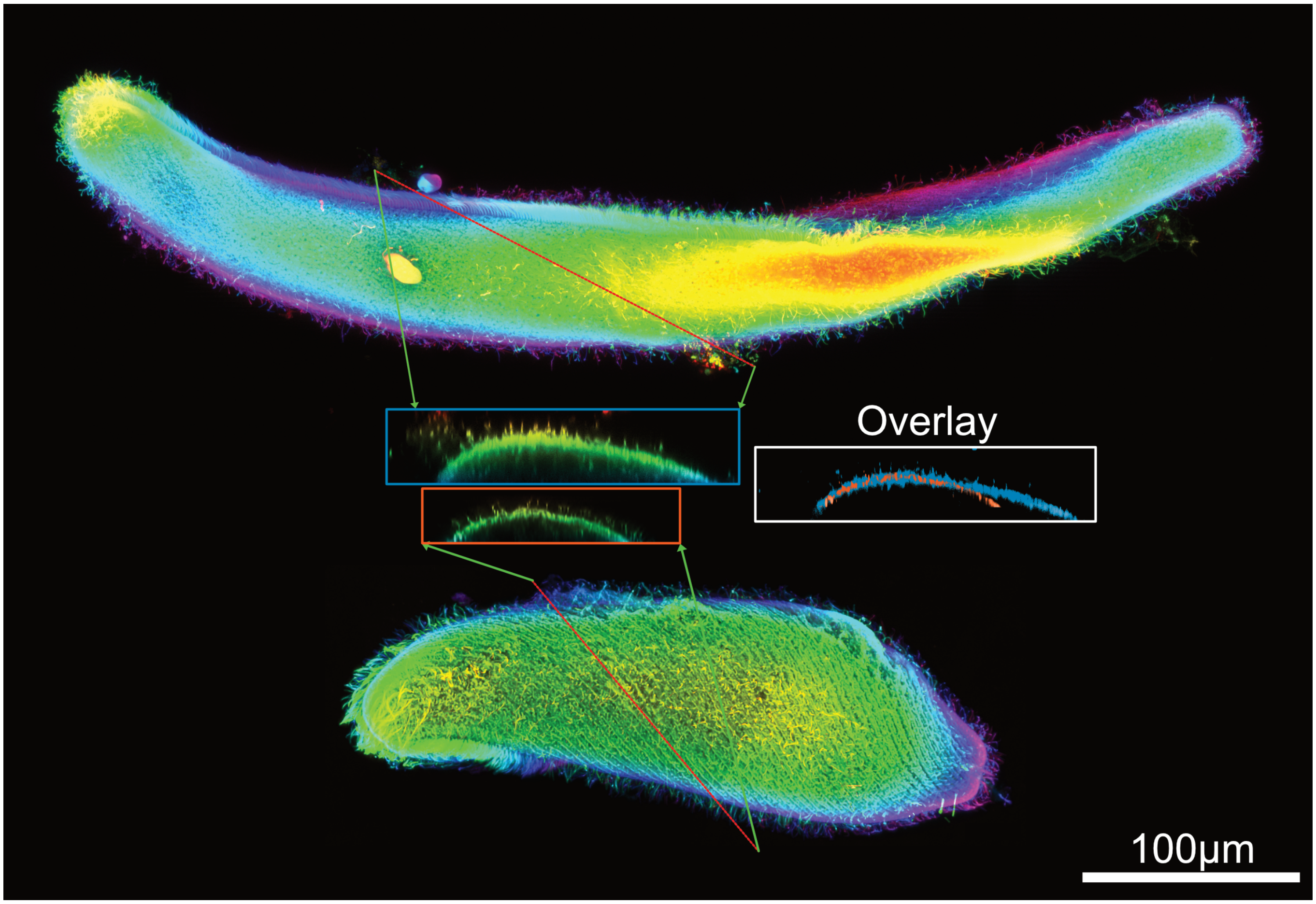


FIG. S1. Comparison of curvature along a furrow of the cortex. Cellmask orange, recolored for depth using ZstackDepthColorCode v.0.0.2 plugin for FIJI (https://github.com/UU-cellbiology/ZstackDepthColorCode). Re-projected using ImageJ along a single ciliary row. Scale bar is 100 µm. Right inset shows overlayed of contracted (orange) and elongated (blue).

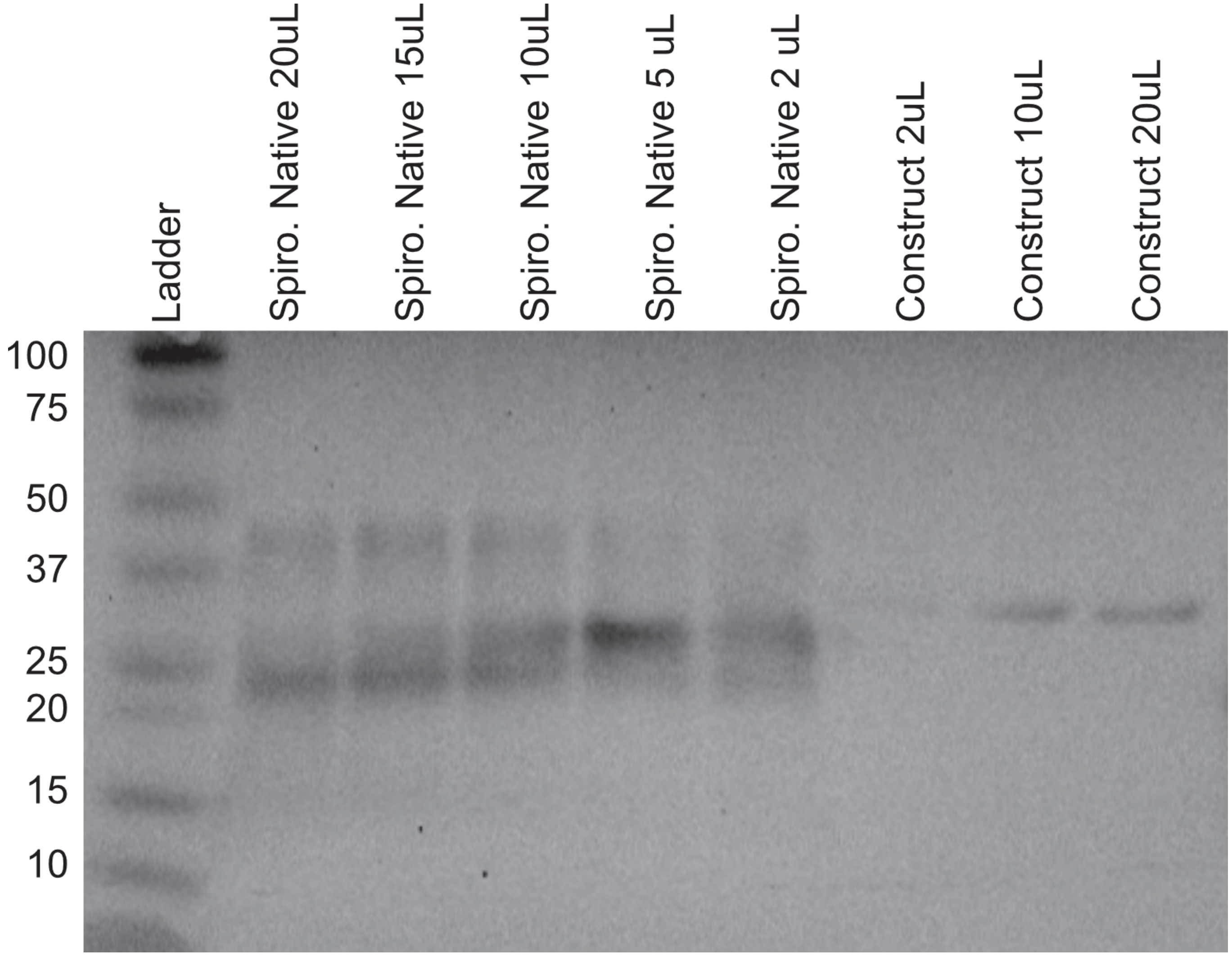


FIG. S2. Western blot of 20H5 antibody against Spirostomum native and recombinant centrin.

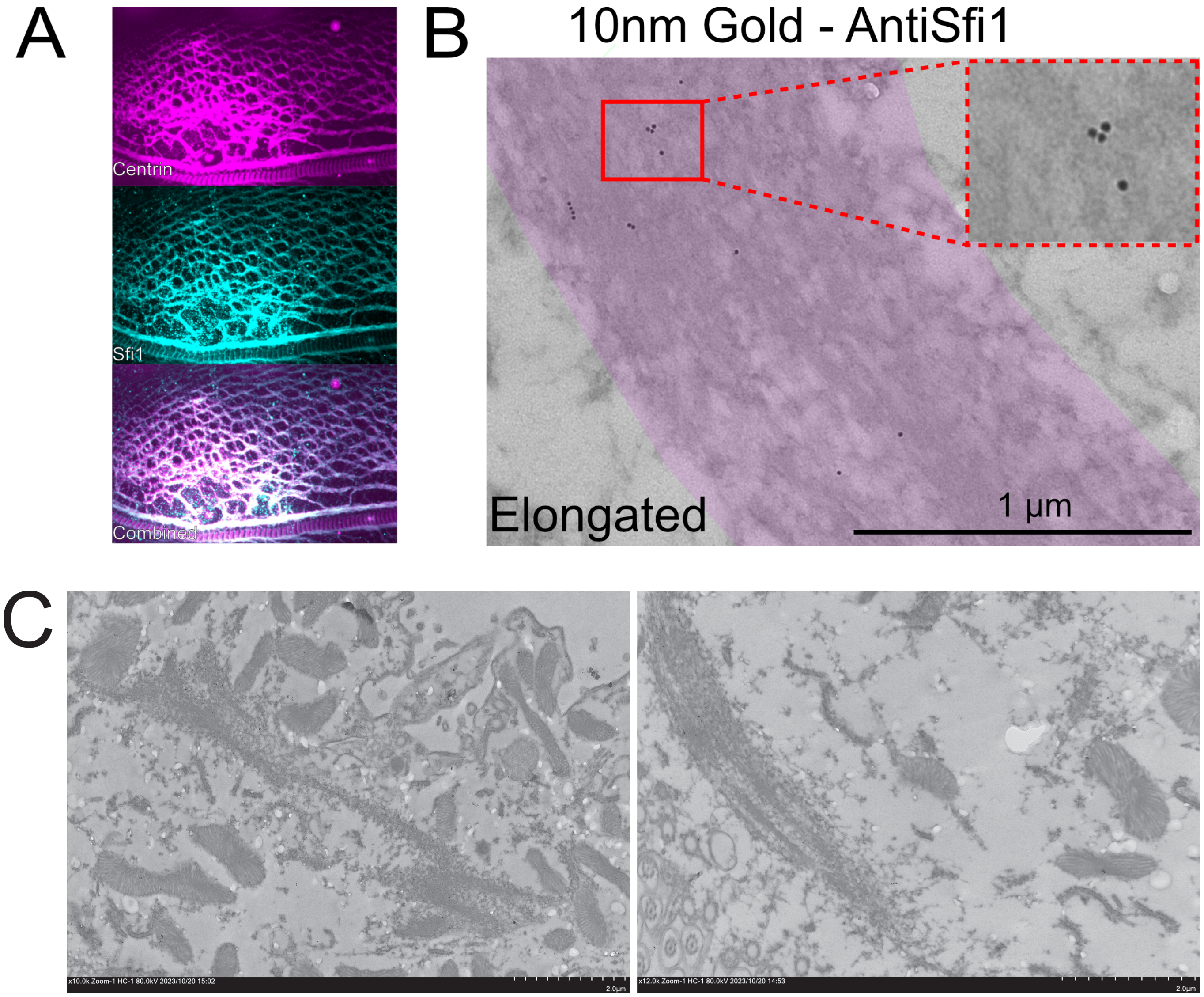


FIG. S3. Localization of Sfi1 in fluorescence microscopy and immunogold TEM. **A** Colocalization of Sfi1 and Centrin in elongated *Spirostomum* Top: Centrin (20H5) is shown in magenta, Middle: Sfi1 (custom peptide antibody against RTEKL-RNALNRVPR), Bottom: combined image. Centrin and Sfi1 exhibit consistent co-localization across the myoneme mesh, but there are distinct bands of centrin present in the oral apparatus that do not fully co-localize with Sfi1. **B** 10nm gold Anti-Sfi1 in elongated Spirostomum. **C** Additional images of Sfi1 labeled TEM, Right is zoom out of B.

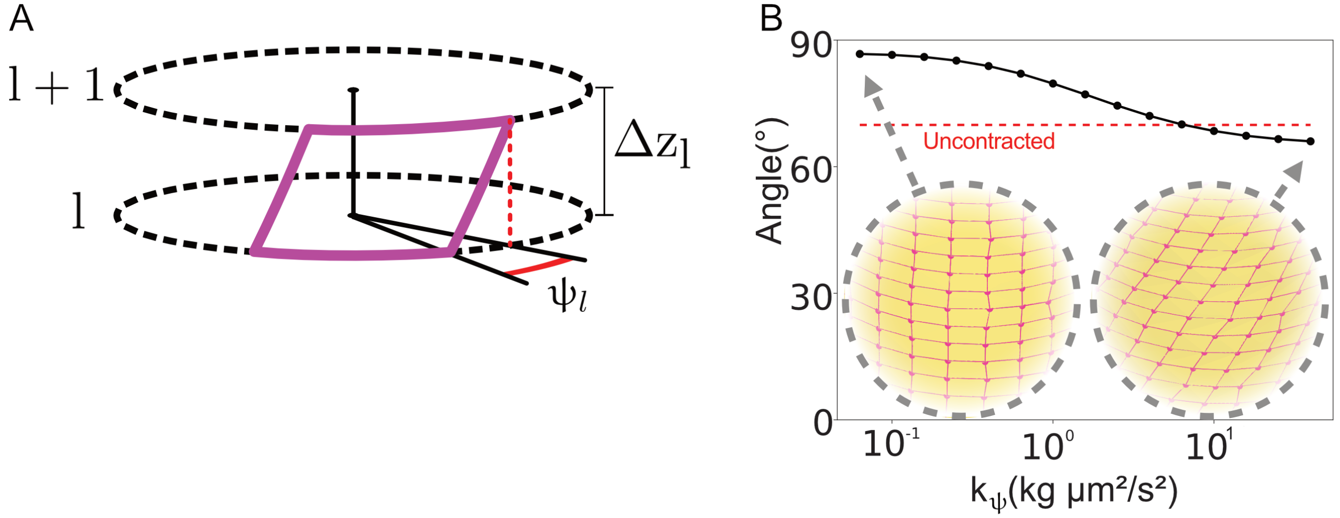


FIG. S4. Graphical Depiction of angle ψ and changes in ϕ with force applied in the latitudinal mesh model. **A** ψ is the angle measured between layers of the mesh as depicted here. **B** Contracted helix angle $\phi_C$ as the angular stiffness $k_\psi$ is varied. The angle of the *elongated* helix $\phi_E$ is shown for comparison in red. Insets show mesh structures for example values of $k_\psi$.

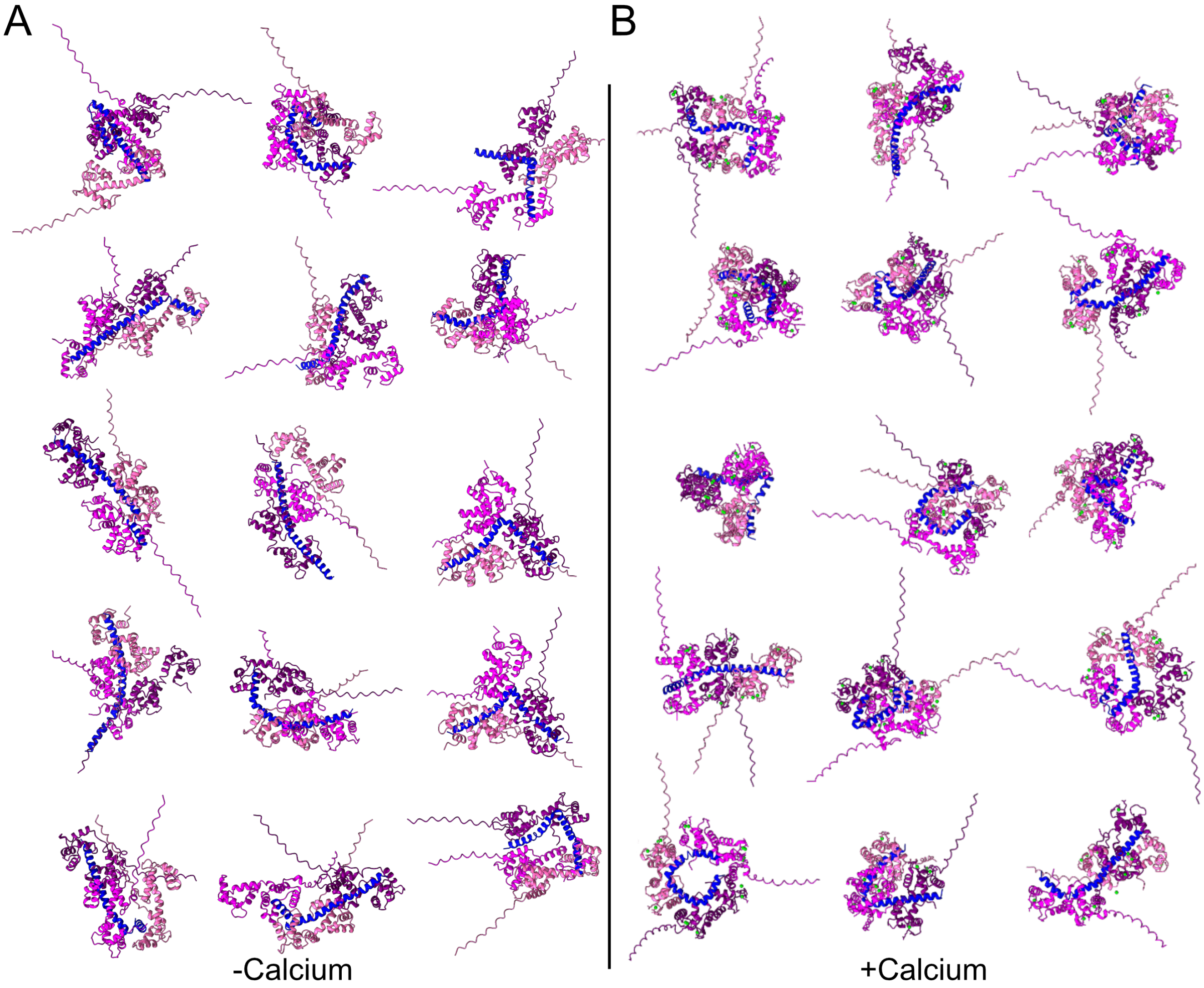


FIG. S5. Additional AlphaFold3 predictions of centrin-Sfi1 complex with and without calcium ions. Alphafold modeling of one sfi1 repeat (blue), with 3 centrin molecules (magenta hues), with and without 12 additional calcium ions (green). **A** Predictions without calcium. **B** Predictions with 12 calcium ions. Models are aligned to best show the conformation of the Sfi1 repeat. Structure files are provided in Supp. Data 3.

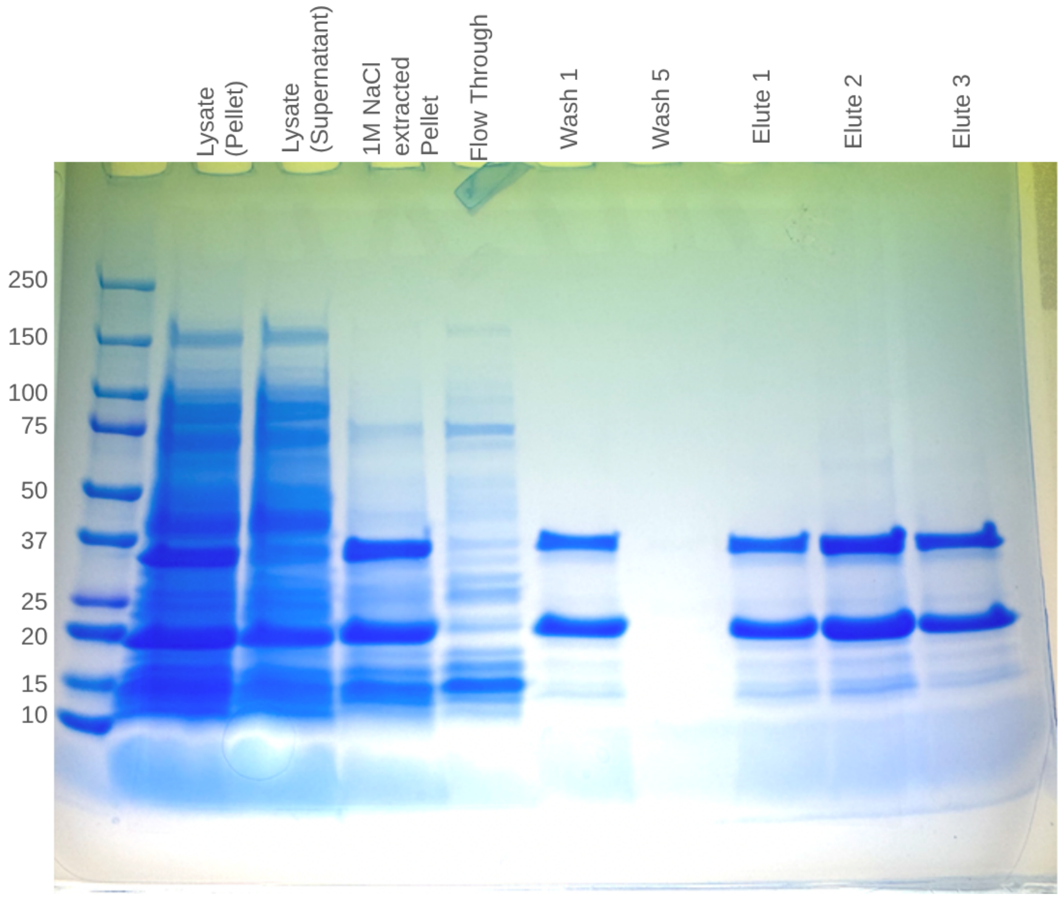


FIG. S6. SDS-Page gel of each preparation stage for the centrin-sfi1x3 construct. Expected mass for the centrin construct is 20.9 kDa and the expected mass for the Sfi1x3 construct is 36.6 kDa.

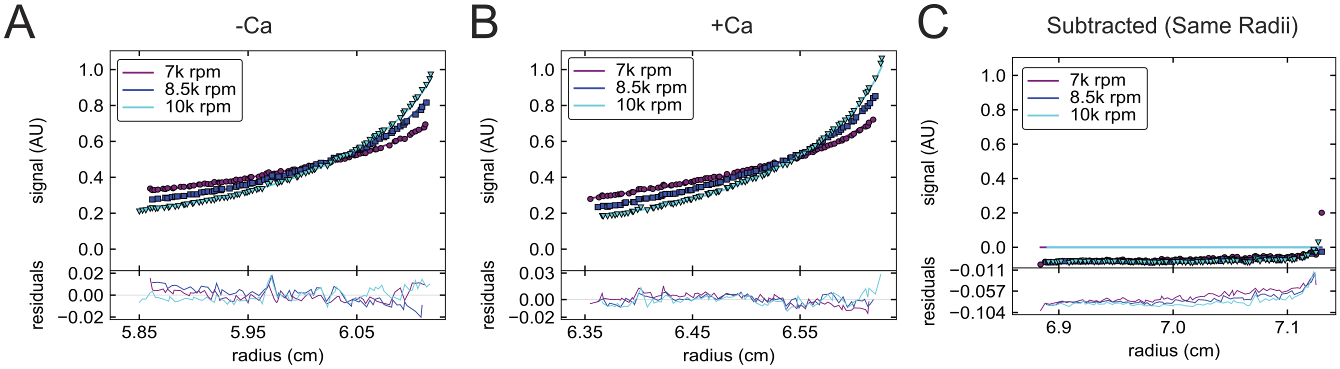


FIG. S7. Steady state AUC measurements of the 3xSfi1 repeat and centrin construct **A** Cen/Sfi1 in 1M NaCl, 10mM EGTA shown at 3 different velocities. Measured mass in EGTA was $121.8 \pm 4.0$ kDa (85%) and $830.5 \pm 424.5 kDa$ (15%) 95% confidence interval. **B** Cen/Sfi1 in 1M NaCl, 10mM EGTA shown at 3 different velocities. Measured mass in Calcium was $121.4 \pm 3.0$ kDa (87%) and $901.7 \pm 247.5$ kDa (13%) 95% confidence interval. **C** Previous two conditions subtracted from each other with the -Ca condition acting as a blank to show a direct comparison between the two conditions. A slight difference is seen at the very end where the highest molecular mass species would be.

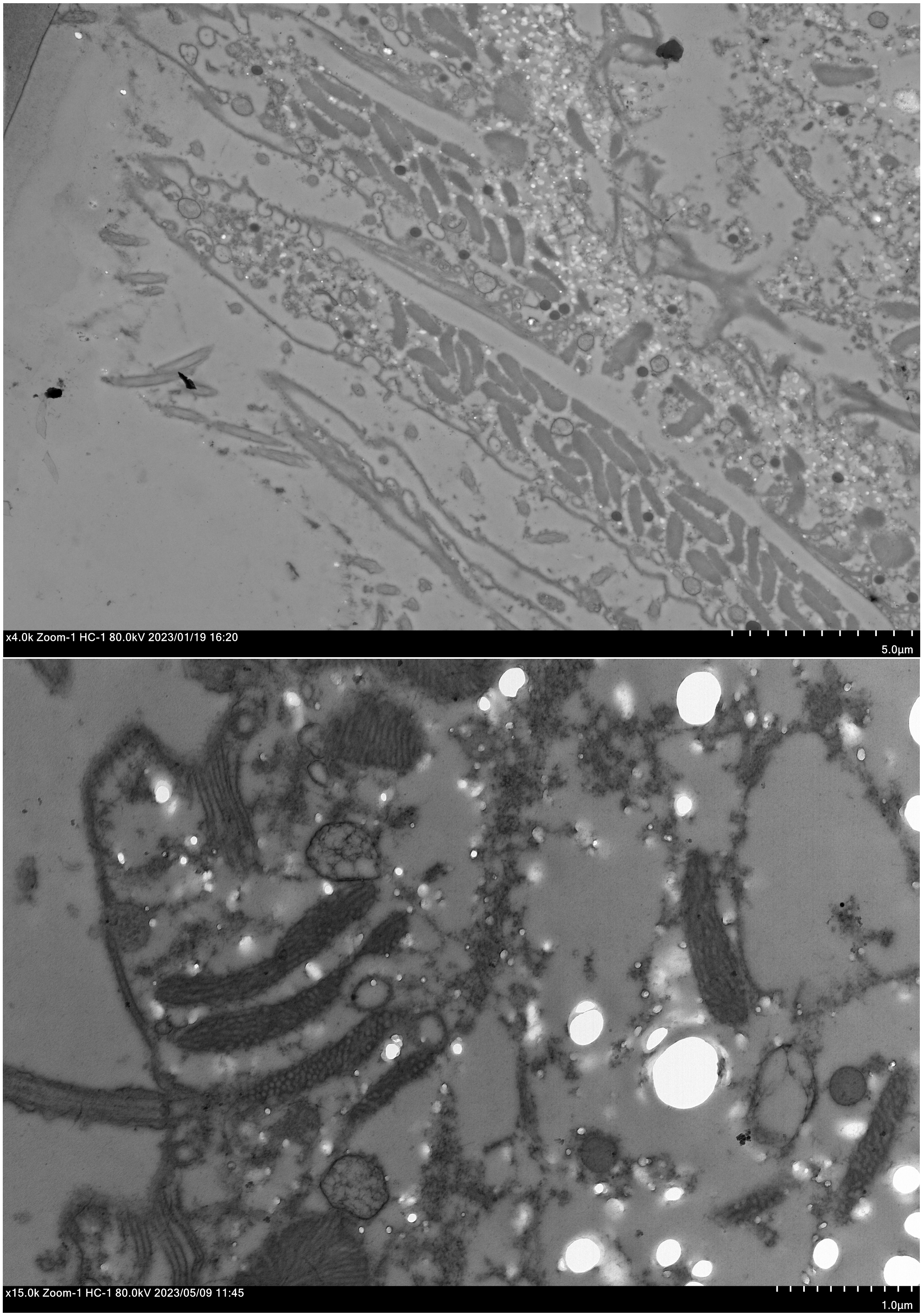


A

B

FIG. S8. Secondary antibody control for immuno TEM imaging
**A** Secondary antibody for 10nm gold anti-mouse. **B** Secondary antibody for 10nm gold anti-rabbit.

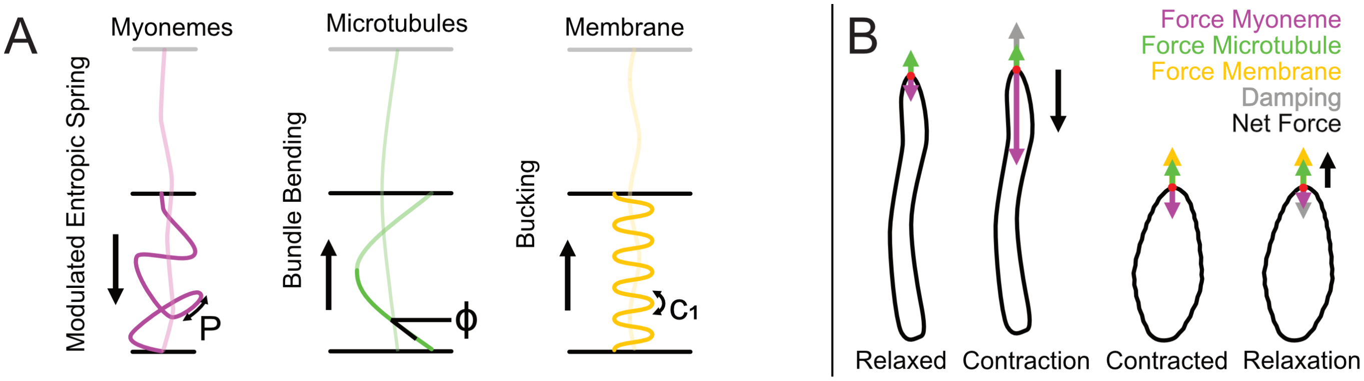


FIG. S9. Overview of forces estimated in the *Spirostomum* cortex. **A** Cartoon of forces estimated (left to right): myoneme (magenta), estimated as an entropic spring; microtubules (green) estimated via their bending moduli; and membrane curvature (gold) estimated via buckling force. **B** Cartoon of the relative magnitudes and net forces acting on *Spirostomum* over a contraction cycle. In the relaxed state, we predict that relatively little force is generated by the myoneme, which balances with the relatively small forces we estimate are contributed by the microtubules and membrane. During contraction, the myoneme generates large forces opposed primarily by drag (damping). In the fully contracted state, the organism is again briefly at equilibrium, when we predict that the myoneme generates relatively small forces compared to those it generates during the contraction phase. During relaxation, we predict that the net force likely requires unknown contributors, since we estimate that the forces from bending the microtubules and membrane are relatively small. Note arrows are not to scale, and only indicate qualitatively relative magnitude.

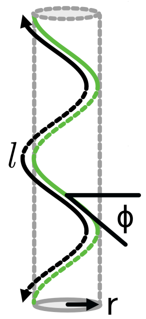


FIG. S10. Parameters for calculating microtubule bending energy. The microtubule bundles (green) form a helix around the cortex of *Spirostomum*, here approximated as a cylinder. Figure indicates the helical angle of the microtubule bundle ϕ, the radius $r$, and the total length of the microtubule bundle $l$ .

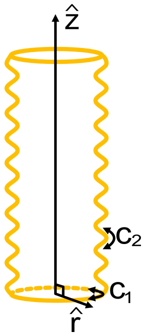


FIG. S11. Parameters for calculating membrane buckling energy. The membrane is approximated as a cylinder with sinusoidal walls. The z-axis (through the middle of the cylinder) and the r-axis (radial to cylinder) are shown. Curvature $c_1$ is in the plane perpendicular to $\hat{z}$ and is $1/r$ where $r$ is the average radius of the cylinder. Curvature $c_2$ is in the $\hat{r}$ plane, along the sinusoidal ridges, and is estimated as the curvature of the sin function.

### SUPPLEMENTAL TABLES

TABLE S1. Summary of Significant Peptide Alignments (Query: RTEKLRNALNRVPR against GSBP2 [1])

| Match | Sbjct Range | Query Aligned | Score (bits) | E-value | Identities | Positives | Gaps |
|---|---|---|---|---|---|---|---|
| **Match 1** | 842 to 850 | 9/9 (**RTEKLRNAL**) | 31.2 | 1e-05 | 9/9 (100%) | 9/9 (100%) | 0/9 (0%) |
| **Match 2** | 7481 to 7490 | 10/14 (**LRNALNRVPR**) | 27.4 | 3e-04 | 8/10 (80%) | 9/10 (90%) | 0/10 (0%) |
| **Match 3** | 10308 to 10317 | 10/14 (**LRNALNRVPR**) | 27.4 | 3e-04 | 8/10 (80%) | 9/10 (90%) | 0/10 (0%) |
| **Match 4** | 8679 to 8691 | 13/14 (**RTEKLRNALNRVP**) | 23.5 | 0.006 | 8/13 (62%) | 9/13 (69%) | 0/13 (0%) |

| **Parameter** | **Symbol** | **Value** |
|---|---|---|
| Structure radius | $R$ | 50 μm |
| Structure length | $L$ | 980 μm |
| Elongated helix angle | $\phi_E$ | 58° |
| Myoneme stiffness per length | $EA$ | 0.01 kg μm/s$^2$ |
| Equatorial shrinking factor (LM) | $\gamma_e$ | 0.8 |
| Longitudinal shrinking factor (LM) | $\gamma_l$ | 0.4 |
| Left-handed shrinking factor (FM) | $\gamma_{l.h.}$ | 0.35 |
| Right-handed shrinking factor (FM) | $\gamma_{r.h.}$ | 0.35 |
| Twisting stiffness (LM) | $k_\psi$ | 2.0 kg μm$^2$/s$^2$ |
| Twisting stiffness (FM) | $k_\psi$ | 0.0 kg μm$^2$/s$^2$ |
| Volume conservation stiffness | $k_V$ | $10^9$ kg/μm s$^2$ |

TABLE S2. Default parameters for the mesh model LM: Latitudinal mesh, FM: fishnet mesh

| ID | Prob | P-val | Loc | Sequence |
|---|---|---|---|---|
| A1 | 98.76 | 2.5e-23 | 1-51 | ------------------PKSALAKWRQFVDDIKKGRILNAVKAQKLATSLTKIPVRKLKDANDRIIGG |
| A2 | 99.86 | 1.5e-28 | 52-120 | GSKVKGVLKGIMKKLSDKPKSAFAKWRKFVDDIKKGKVLDAVKAQKLAVCLAKIPVRRLKDANDRIIGG |
| A3 | 99.25 | 3.1e-24 | 121-189 | GSKIKGVLRGIMKKLSERPKSAFAKWRAYIDSVKKGKILDAVKAQKLAASLRKIPVRRLKDANDRILGG |
| A4 | 99.71 | 7.4e-27 | 190-258 | GSKVAGVLRGIMKKLKDKPKSALAKWRDFVDNIKKGKILNAVKAQKLAVCLAKIPVRRLKDANDRIIGG |
| A5 | 99.62 | 3.0e-26 | 259-327 | GSKIAGVLRGIMKKLKDKPKSAFAKWRQFIEDIKKGKVLDAVKAQKLASCLTKIPVRKLKDANDRIIGA |
| A6 | 99.67 | 1.5e-26 | 328-396 | GSKVAGVLRGIMKKLKDKPKCALARWRQYVDEVKKGKVLDAVKAQKLAASLNRIPVRIMKDANDRILGD |
| A7 | 99.60 | 4.7e-26 | 397-465 | GSKIKGVLRGIMKKLAEKPKKAFARWRAFVDDIKKGKVLDAVKAQKLAACLAKIPVRRLKDANDRILGG |
| A8 | 99.52 | 1.4e-25 | 466-534 | GSRIAGALRGIMKKLKDKPKSALARWRQFVEDIKKGKVLDAVKAQRLAASLSKIPTRRLKDANDRIIGG |
| A9 | 99.34 | 9.8e-25 | 535-603 | GSKVAGVLRSIMSRLKEKPKSAFTKWRAFVNDIKKGRMLNAVKAQQLVVSLSKIPIRRLKDANDRIIGG |
| A10 | 99.70 | 8.4e-27 | 604-672 | GSKIKGVLRAILKKLKDKPKSALAKWRTFVDDIKKGKVLDAVKAQKLAACLARIPVRRLKDANDRVIGG |
| A11 | 99.78 | 1.5e-27 | 673-741 | GSKIKGVLRSIMKKLSDKPKSAIARWRQFVDDIKKGKVLDAVKAQKLASCLARIPNRIVRDANDRILGD |
| A12 | 99.10 | 9.8e-24 | 742-810 | GSKVKGALRALVKKLSDKPKTALKKWKQFVDDVKKGKVLDAVKAQKLVVALSRVPIRIVKDANERILGG |
| A13 | 99.64 | 2.9e-26 | 811-879 | GNKVKGALRSLVHKLAQKPKSALSKWRQFVEDVKKGKVLDAVKAQKLASTLSRVPARVVRDANDRILGG |
| A14 | 99.57 | 7.0e-26 | 880-948 | GDKVKGALRALVHKLSQKPKLALSKWRSFVEGVKKGRILDAVKAQKLAKSLSRVPLRTVKDANDRVIGG |
| A15 | 99.67 | 1.4e-26 | 949-1017 | GNKVKGALRNLMHKLIQKPRQALEKWKQYVENVKRGKILDQVKVQKLKASLIRVVTRTLFPAFKAATGM |
| A16 | 97.73 | 3.4e-19 | 1018-1068 | PTLIKSMLRNLAHIIDATPRNALSRWRSAIASSKHNAAQESLKNVLRAIKF------------------ |

TABLE S3. *Spirostomum* Sfi1 15 repeat self-alignment. Conserved tryptophans shown in green. Helix breaking residues (proline and glycine) shown in blue. Probabilities and p-values for the alignment were calculated using MPI Bioinformatics Toolkit (see Supp. Materials and Methods).

| ID | Prob | P-val | Loc | Sequence |
|---|---|---|---|---|
| A1 | 82.33 | 7.6e-08 | 12-37 | ---SFRNTWLLFRSFQ.QWITLTQTLKEQS............ |
| A2 | 80.10 | 1.2e-06 | 38-58 | ---RLADQAFLNKMFR.KILKAQEH-----WKHLET....... |
| A3 | 84.69 | 5.0e-08 | 65-87 | VNTDNIKKIFLRTTFH.IWKLRHK------EINY......... |
| A4 | 72.88 | 1.0e-05 | 92-108 | -------HGLERRIFE.RIKQKVIN-----YEYNKS....... |
| A5 | 84.82 | 6.3e-08 | 115-138 | IAEKVRSFSLQRKYLN.KWEKKNIE-----NEDKLG....... |
| A6 | 83.77 | 3.9e-07 | 145-168 | ALYELENKFIKQKFFR.KLNRSFQH-----SQQEAIAKS.... |
| A7 | 80.63 | 1.3e-06 | 178-198 | ----KLNQTLLRCVFEKMWLKRFED-----HLHLYS....... |
| A8 | 81.62 | 1.3e-06 | 205-232 | -IVSLKEANLVKRIFH.SWKKLLYIDLKAS............ |
| A9 | 82.30 | 7.1e-08 | 233-253 | ---DYSRTNLLKSSLR.SWKLEVKL-----KIFEQ........ |
| A10 | 79.51 | 2.9e-06 | 259-283 | ----KCKKSIQASAYR.TWRKRIQYGKISS............ |
| A11 | 80.86 | 3.1e-06 | 284-304 | ---EHVKTAFCAKYLG.VWKRRMLQ-----MNSMND....... |
| A12 | 83.58 | 3.1e-07 | 311-339 | EASKFYEEGLVNECLA.IWKERLIKTKELE............ |
| A13 | 76.34 | 7.4e-05 | 340-360 | ---DRYNFLCKTHAIL.TVKRTLMH-----IDNVHLLYTKLAP |
| A14 | 79.77 | 2.4e-06 | 374-393 | ----SMDRVKLSKAFL.KWRKATRF-----KVRHKLNDILHVY |
| A15 | 81.82 | 7.5e-07 | 407-427 | --EKSKERELQSQLFN.AWRNRFC------............ |

TABLE S4. *S. cerevisiae* Sfi1 15 repeat K246-E677 self-alignment [2]. Loc 1 is residue K246. Conserved tryptophans shown in green. Helix breaking residues (proline and glycine) shown in blue. Probabilities and p-values for the alignment were calculated using MPI Bioinformatics Toolkit (see Supp. Materials and Methods).

| Construct | Vector | DNA Sequence inserted | Translated AA Sequence |
|---|---|---|---|
| Centrin | pj441 | AAGGAGGTAAAAAATGCATCATCATCACCATCACAT<br>GGCAGGCAGAGCACCAGCATCGAAGGCGGGTCCACC<br>GGCGGCGAAAGCTGGCCCGCCGGCGTTCAACGCGAA<br>GCGTTATGAGCGCCCTGGTCTGACCGAAGATGAAAT<br>CGAAGAGATCAAAGAGGCCTTTGACCTGTTTGATAC<br>TGACGGTAGCGGTACCATTGACCCGAAAGAACTGAA<br>GTCCGCGATGGAAAGCCTGGGCTTTGAGGCAAAGAA<br>CCAGACGATTTACCAGATGATTTCTGATCTGGACAA<br>GGACGGCAGCGGTGCCATTGACTTCGATGAATTCCT<br>GGACATGATGACCGCGCGTCTGAGCGATAAAGACAG<br>CCGCGACGACATCAATAAAGTGTTTCGTCTGTTCGA<br>CGATGAGAAGCAAGGTTTTATCACGATTAAGAACTT<br>GCGCCGTGTCGCCAAAGAACTGGGTGAAACCATGAC<br>CGATGAAGAACTGTTGGAAATGATCGAGCGTGCAGA<br>TAGCGACGGCGATGGTCGTGTTACCGCGGAAGATTT<br>CTACAATATCATGACGAAAAAAGCTTTCCCGTAATA<br>A | MHHHHHHMAGRAPASKAGPPAAKAGPPAFNAKRYER<br>PGLTEDEIEEIKEAFDLFDTDGSGTIDPKELKSAME<br>SLGFEAKNQTIYQMISDLDKDGSGAIDFDEFLDMMT<br>ARLSDKDSRDDINKVFRLFDDEKQGFITIKNLRRVA<br>KELGETMTDEELLEMIERADSDGDGRVTAEDFYNIM<br>TKKAFP* |
| 3xSfi1-SH3 | pj414 | AAGGAGGTAAAAAATGCACCATCACCATCATCACGC<br>GGAATACGTTCGTGCTTTGTTCGACTTTAACGGCAA<br>TGATGAAGAAGATTTGCCGTTTAAGAAAGGTGACAT<br>CCTGCGTATCCGCGATAAGCCTGAAGAACAGTGGTG<br>GAATGCTGAGGACTCTGAGGGTAAGCGTGGTATGAT<br>TCCGGTGCCGTATGTGGAGAAGTATCGCGCAGAGGC<br>TGCAGCCAAAGAGGCAGCGGCCAAAGCGCCGAAAAG<br>CGCGTTCGCCAAGTGGCGTGCGTACATCGATAGCGT<br>GAAGAAAGGTAAGATTCTGGATGCAGTCAAGGCCCA<br>AAAACTTGCGGCGTCCCTGCGTAAGATTCCGGTACG<br>CCGTCTGAAAGACGCCAACGACCGCATCCTGGGTGG<br>CGGCTCCAAAGTCGCGGGTGTGCTGCGCGGTATTAT<br>GAAAAAGCTGAAAGATAAGCCGAAATCGGCACTGGC<br>GAAGTGGCGTGACTTCGTTGATAACATCAAGAAAGG<br>CAAGATTCTGAACGCCGTCAAAGCCCAGAAACTGGC<br>TGTGTGTTTAGCGAAGATCCCGGTTCGTCGCCTGAA<br>AGATGCGAATGACCGCATCATCGGTGGCGGCAGCAA<br>GATTGCTGGTGTGTTGCGTGGTATCATGAAAAAACT<br>GAAAGACAAACCGAAGTGCGCCCTGGCGCGTTGGCG<br>CCAATACGTCGACGAAGTCAAAAAGGGTAAGGTTCT<br>GGACGCAGTTAAAGCGCAAAAACTGGCGGCGAGCCT<br>GAACCGTATTCCGGTTCGCATTATGAAAGACGCTAA<br>TGACCGTATCCTGGGTGATGGTAGCAAGATTAAAGG<br>CGTGCTGCGTGGCATTATGAAGAAACTGGCAGAGAA<br>GCCAGCCGAGGCAGCAGCGAAGGAAGCGGCAGCGAA<br>AGCACCGCCACCGGCACTGCCGCCGAAGCGTCGTAG<br>ACCGCCGCCGGCGTTGCCTCCGAAGCGCCGTCGTGA<br>GCAGAAACTGATTAGCGAAGAAGATCTCTAATAA | MHHHHHHAEYVRALFDFNGNDEEDLPFKKGDILRIR<br>DKPEEQWWNAEDSEGKRGMIPVPYVEKYRAEAAAKE<br>AAAKAPKSAFAKWRAYIDSVKKGKILDAVKAQKLAA<br>SLRKIPVRRLKDANDRILGGGSKVAGVLRGIMKKLK<br>DKPKSALAKWRDFVDNIKKGKILNAVKAQKLAVCLA<br>KIPVRRLKDANDRIIGGGSKIAGVLRGIMKKLKDKP<br>KCALARWRQYVDEVKKGKVLDAVKAQKLAASLNRIP<br>VRIMKDANDRILGDGSKIKGVLRGIMKKLAEKPAEA<br>AAKEAAAKAPPPALPPKRRRPPPALPPKRRREQKLI<br>SEEDL* |

TABLE S5. Constructs used in *in vitro* reconstitution of myonemal proteins.

| Construct | Vector | DNA Sequence inserted | Translated AA Sequence |
|---|---|---|---|
| SH3-SpectrinControl | pET-21a(+) | CCCCCCCCAGCTCTACCGCCTAAAAGAAGGCGTCCG<br>CCACCGGCGCTCCCGCCTAAAAGACGTCGCATGGTT<br>CATCAGTTCTTCCGCGATATGGATGATGAAGAGTCT<br>TGGATCAAAGAAAAAAAGCTGCTGGTGTCCAGCGAA<br>GACTATGGCCGTGACTTGACCGGTGTTCAGAATCTG<br>CGTAAAAAGCATAAACGCTTGGAGGCCGAACTGGCT<br>GCGCATGAACCGGCGATCCAAAGCGTGCTGGATACT<br>GGTAAGAAACTGTCTGATGACAACACGATTGGTAAG<br>GAGGAAATCCAACAACGCCTCGCGCAATTCGTCGAT<br>CATTGGAAAGAGCTGAAGCAGCTGGCGGCTGCGCGT<br>GGTCAGCGTCTTGAAGAGTCGCTGGAATACCAGCAA<br>TTCGTGGCCAATGTTGAAGAGGAGGAGGCGTGGATC<br>AACGAGAAGATGACCCTGGTCGCGAGCGAAGACTAC<br>GGCGACACGCTGGCAGCGATCCAGGGTTTGCTGAAA<br>AAGCACGAGGCCTTTGAAACCGACTTCACCGTTCAC<br>AAAGATCGCGTGAATGACGTTTGCGCGAACGGTGAA<br>GATTTGATCAAAAAGAACAACCATCACGTGGAAAAT<br>ATTACCGCAAAAATGAAGGGCCTGAAGGGCAAAGTT<br>TCCGACCTGGAGAAGGCTGCTGCCCAACGCAAGGCG<br>AAGCTGGACGAGAACAGCGCAGCGGAATACGTGCGT<br>GCACTGTTTGATTTTAACGGAAACGATGAAGAGGAT<br>TTGCCGTTTAAAAAGGGCGACATCCTGCGTATTCGT<br>GACAAACCGGAAGAGCAGTGGTGGAATGCAGAGGAC<br>AGCGAAGGTAAGCGTGGCATGATTCCGGTTCCGTAT<br>GTAGAGAAGTATCGTGAACAGAAATTGATTAGCGAG<br>GAGGACTTACATCACCACCACCACCAC | PPPALPPKRRRPPPALPPKRRRMVHQFFRDMDDEES<br>WIKEKKLLVSSEDYGRDLTGVQNLRKKHKRLEAELA<br>AHEPAIQSVLDTGKKLSDDNTIGKEEIQQRLAQFVD<br>HWKELKQLAAARGQRLEESLEYQQFVANVEEEEAWI<br>NEKMTLASEDYGDTLAAIQGLLKKHEAFETDFTVHK<br>DRVNDVCANGEDLIKKNNHHVENITAKMKGLKGKVS<br>DLEKAAAQRKAKLDENSAAEYVRALFDFNGNDEEDL<br>PFKKGDILRIRDKPEEQWWNAEDSEGKRGMIPVPYV<br>EKYREQKLISEEDLHHHHHH |

TABLE S6. Constructs used in *in vitro* reconstitution of myonemal proteins (Continued).

### SUPPLEMENTAL DATA

The following files are included as data referenced in this study.

| | File Name | Description |
|---|---|---|
| 1 | `Yeast_15rp_AF2.pdb` | Representative AlphaFold2 prediction for 15 *S. cerevisiae* Sfi1 repeat (equivalent to 5 *Spirostomum ambiguum*). |
| 2 | `Spiro_1rp_AF2.pdb` | Representative AlphaFold2 prediction for 1 *Spirostomum ambiguum* Sfi1 repeat. |
| 3 | `Spiro_1rpSfi1_3cen_+-Ca_30runs.pdb` | Additional AlphaFold3 predictions of 1 Sfi1 repeat with 3 centrins 30 total complexes. |

*Spirostomum ambiguum* Centrin Sequence

```
MAGRAPASKAGPPAAKAGPPAFNAKRYERPGLTEDEIEEIKEAFDLFDTDGSGTIDPKELKSAMESLGFEAKNQTIYQMISDLDKDGSGAIDFDEFL
DMMTARLSDKDSRDDINKVFRLFDDEKQGFITIKNLRRVAKELGETMTDEELLEMIERADSDGDGRVTAEDFYNIMTKKAFP
```

### SUPPLEMENTAL DISCUSSION

#### 1. Energetics and total force calculations

In this section, we estimate the mechanical energy stored in and forces exerted by structures in the *Spirostomum* cortex (Fig. S9). Starting with our imaging results shown in the main text and previously measured mechanical parameters, we estimate energies stored in cortical microtubule bundles and the membrane and use these estimates to make predictions about the force exerted by the myonemes.

*Microtubule bending energy estimation*

As shown in Fig. 1, the microtubules in the cortex of *Spirostomum* form large helical coils, similar in shape to a spring. During contraction, these coils compress, which could potentially result in the storage of energy from contraction in microtubule bending. We used previous measurements of the mechanical properties of microtubules to estimate the total energy stored in this bending. A comparison of the curvature of the microtubules can be seen in Fig. S1.

The elastic energy stored in a single microtubule curled into a transverse coil (Fig. S10) is given by

$$E = \frac{\kappa l}{2r^2}\cos^4(\phi) \tag{S1}$$

where $E$ is the total elastic energy in the coil, $\kappa$ is the bending stiffness of the microtubule, $l$ is the total length of the microtubule, $r$ is the radius of the coil, and $\phi$ is the complement of the pitch angle of the coil (we have reframed Eqn. B2 from [3] to match the angle as we have defined it in Fig. 1A). For a bundle of microtubules, we expect this energy to scale linearly with the number of microtubules in the bundle.

In the calculations below, we use the following estimated parameters (based on imaging data from elongated *Spirostomum* as in Fig. 1 and previously published measurements of microtubule mechanics):

$\kappa \approx 10\,\mathrm{pN\,\mu m^2}$ [3–5]

$r \approx 35\,\mathrm{\mu m}$

$l \approx 900\,\mathrm{\mu m}$

$\phi \approx 64°$

$\approx 150$ microtubules per bundle [6] and $\approx 50$ bundles in the organism.

Plugging in these values, we find

$$E \approx 1.0\,\mathrm{fJ}. \tag{S2}$$

In contracted *Spirostomum*, we estimate the following changes to these parameters (those not listed are unchanged):

$r \approx 60\,\mathrm{\mu m}$

$\phi \approx 34°$

Plugging in these values, we find

$$E \approx 4.4\,\mathrm{fJ}. \tag{S3}$$

Per microtubule this works out to be $\sim 1 \times 10^{-4}\,\mathrm{fJ}$. Notably, since the radius of the organism expands during contraction while the helical pitch decreases, there is relatively little resulting change in energy stored in microtubule bending. Compared to the amount of energy expended to cause organismal contraction (see below), we thus expect microtubules to have a negligible contribution toward opposing contraction.

*Membrane buckling energy estimation*

In the contracted state, we observe periodic ripples in the membrane (Fig. 1). To measure the potential effect of membrane buckling during contraction and to determine if the membrane stores significant energy from contraction for use in elongation, we estimate the energy expended into the membrane to form these ridges.

The energy of bending for a membrane is given by

$$E_m = \iint \frac{1}{2} K_b (c_1 + c_2 - c_0)^2 + \bar{K} c_1 c_2 \, dA \tag{S4}$$

where $c_1$ and $c_2$ are the principal curvatures, and $c_0$ is the spontaneous curvature, $K_b$ is the bending rigidity and $\bar{K}$ is the Gaussian bending modulus (Helfrich theory [7, 8]). We approximate *Spirostomum* as a wavy cylinder with a sinusoidal curve that varies along the long axis of the cylinder (Fig. S11). The curvature $c_1$ is defined by the radius of this cylinder, whereas $c_2$ is caused by the membrane ridges. We are actually interested in the *change* in bending energy during contraction, and assume that it is dominated by $c_2$, which in the contracted state is much larger than $c_1$. We also assume that, while the spontaneous curvature $c_0$ may be non-zero, it is still much smaller than $c_2$. However, we acknowledge that there may be some mechanisms for manipulating spontaneous curvature that are yet unknown in *Spirostomum*, and that such mechanisms might be useful in inducing the changes in shape we observe.

Thus, we now want to evaluate

$$\Delta E_m \approx \iint \frac{1}{2} K_b c_{2,c}^2 \, dA \tag{S5}$$

where the subscript $c$ indicates the contracted state. To find $c_{2,c}$, we consider the surface described above, which is defined by

$$r = r_0 + \alpha \sin(\omega z) \tag{S6}$$

where $\alpha$ is the amplitude of the sinusoidal wave, $\omega$ is its angular frequency, and $r_0$ is the average radius of the organism. The curvature of this function is

$$c_{2,c} = \frac{\alpha w^2 \sin(wz)}{(1 + \alpha^2 \omega^2 \cos^2(wz))^{\frac{3}{2}}}. \tag{S7}$$

We now take advantage of the periodicity of this function by evaluating the integral in $z$ over a single period and scaling it up to the entire organism:

$$\Delta E_m \approx \frac{1}{2} K_b \iint c_{2,c}^2 \, dA = K_b r L \frac{\omega}{2} \int_0^{2\frac{\pi}{\omega}} c_{2,c}^2 dz \tag{S8}$$

where $L$ is the length of the organism and $r$ is its radius. This integral evaluates to:

$$\frac{\alpha^2 w^4 (4 + 3\alpha^2 \omega^2)}{8(1 + \alpha^2 \omega^2)^{\frac{3}{2}}} \tag{S9}$$

resulting in

$$\Delta E_m \approx \frac{1}{2} K_b \Big( \frac{\alpha^2 \omega^4 (4 + 3\alpha^2 \omega^2)}{8(1 + \alpha^2 \omega^2)^{\frac{3}{2}}} \Big) \cdot 2\pi r L \tag{S10}$$

To estimate the value of this expression, we use the following parameters (from previous measurements and imaging as in Fig. 1):

$K_b \approx 1 \times 10^{-19}$ J [9]

$r_c \approx 50\,\mu\text{m}$

$\alpha \approx 1\,\mu\text{m}$

$\omega \approx \frac{2\pi}{2\,\mu\text{m}}$

$L \approx 1\,\text{mm}$

$$\Delta E_m \sim 100\,\text{fJ} \tag{S11}$$

While this energy is somewhat larger than what we estimate is stored in microtubule bending, it is still not a significant source of energy storage compared to that expended in contraction.

*Estimate of the force exerted by individual myoneme filament bundles*

We can now compare our estimates of the mechanical contribution of the microtubules and membrane to previous measurements of the force generated during contraction. Using a glass needle to hold *Spirostomum* in an elongated state, Hawkes et al. measured the force generated by *Spirostomum* contraction to be $5 \times 10^5$ pN [10]. This important measurement was conducted in an isometric state, where enough force was exerted to prevent contraction. To get a rough estimate of the energy expended during contraction, we can multiply this force by the change in length, i.e.,

$$E = \int F \cdot dl \approx F \cdot \Delta l.$$

With a change in length of $\sim 1\,\text{mm}$, this results in an estimate of the work done by the myoneme of:

$$E_{myoneme} \approx 1 \times 10^5\,\text{fJ}. \tag{S12}$$

This value is at least 3 orders of magnitude larger than either our estimate of the energy required to bend the membrane into folds (Eq. S11) or our estimate of the bending energy of microtubules in the contracted state (Eq. S3). Thus, we conclude that neither microtubules nor membrane bending likely oppose contraction to a significant degree.

We can also use this measurement of the organismal force to estimate the force generated by individual myoneme bundles. Taking the myoneme as a mesh of springs, some in parallel and some in series, we find

$$F_{\text{total}} = N_{\text{parallel}} \cdot F_{\text{bundle}}$$

where $N_{\text{parallel}}$ is the number of bundles that effectively act in parallel around the circumference of the organism, and $F_{\text{bundle}}$ is the force produced by a single myoneme bundle. While none of the individual myonemes are perfectly in series or in parallel, we can estimate the number of bundles around the circumference (i.e., acting in parallel) to be approximately 100 from Fig. 1.

This then gives us an estimate of the force generated by each bundle

$$F_{\text{bundle}} \approx 1000\,\text{pN}. \tag{S13}$$

*Estimate of myoneme forces if filaments act as entropic springs*

Finally, we estimate whether a reasonable change in the persistence length of the contracted myoneme fibers could produce the force estimated in the previous section. From TEM images of the myoneme, such as Fig. 4A, which show around 10 filaments per linear cross section, we estimate that each myoneme bundle contains $\sim 10^2 = 100$ filaments, all acting roughly in parallel. This results in an estimate that each filament produces $F_{\text{filament}} \approx F_{\text{bundle}}/100 = 10$ pN (using $F_{\text{bundle}}$ in Eq. S13). AlphaFold predictions and AUC experiments (Fig. 5) suggest one potential molecular mechanism by which helix-breaking residues, modulated by centrin-calcium interactions, could effectively modulate the filament persistence length and contribute to contraction. Here, we validate that these proposed dynamics could reasonably generate force of the appropriate magnitude.

To first order, the Marko–Siggia approximation of the force extension curve for a worm-like chain model is

$$F_{\text{filament}}(x) = \frac{k_B T}{P}\left(\frac{1}{4\left(1-\frac{x}{L}\right)^2} - \frac{1}{4} + \frac{x}{L}\right) \tag{S14}$$

where P is the persistence length, $k_b$ is the Boltzmann constant, $T$ is the temperature, $L$ is the total length of the fiber, and $x$ is the extension of the fiber beyond its equilibrium end-to-end distance [11, 12].

The helix-breaking residues in Sfi1 are regularly spaced at separations of $\sim$ 35 amino acids (Fig. 5A and Supplementary Table S3). With 3.6 amino acids per turn of the alpha helix, this results in a spacing of $\approx$ 10 turns of the alpha helix between these residues, or $\approx$ 5 nm of alpha helical length. We therefore use 5 nm as an order of magnitude estimate of the filament persistence length in the contracted state. Plugging these values into Eq. S14, we find that the ratio of the distance from equilibrium $x$ over the polymer length $L$ required to achieve $F_{bundle} = 10$ pN is $x/L = 0.85$ or 85% extension. In other words, filaments would need to have an end-to-end distance of $\approx$ 85% of their total polymer length in order to generate the necessary force.

From TEM skeletonization measurements, we estimated that the elongated myoneme fibers have a persistence length $\sim$100 nm (Fig. 4). This is 20-fold higher than the 5 nm we estimate if each segment between helix breaking residues becomes an effective segment length when contraction is induced (see above). That is, in Fig. 6B, the segments on the left that are straight in the elongated state are effectively already stretched when contraction is induced, leading them toward the contracted state on the right. We might expect, then, that if the elongated fiber is roughly at equilibrium before contraction, each segment of fiber would be effectively stretched due to the 20-fold difference in persistence length when contraction is induced.

#### 2. Latitudinal mesh model

We find that in the latitudinal model, the closed-loop strands of myoneme lead to local pinching around the equators and create inflection points where the structure is no longer convex. In Fig. 2C we show a heatmap of the inflection metric (which is less than 1 for non-convex shapes; see Materials and Methods) as we vary the myoneme spring shrinking factors $\gamma_l$, $\gamma_e$ of longitudinal and equatorial myoneme strands. This figure reveals a critical line in the $\gamma_l, \gamma_e$ space over which the structure loses convexity due to greater shrinkage in the equatorial direction relative to the longitudinal direction. To achieve appreciable contraction in the organism's length, the shrinkage in the equatorial direction must be significantly less than in the longitudinal direction; this condition is likely non-physiological since it would require systemic differences in myoneme structure or biochemistry depending on their angle. Furthermore, we find that in the latitudinal mesh, the adjacent equatorial strands are free to untwist relative to each other and will do so unless explicitly penalized by the torsional penalty (Fig. S4B). Even under stiff torsional rigidity, the helix angle in the contracted equatorial mesh is much larger than those experimentally observed.
